# Deep Immunophenotyping Reveals Endometriosis is Marked by Dysregulation of the Mononuclear Phagocytic System in Endometrium and Peripheral Blood

**DOI:** 10.1101/2022.02.02.478809

**Authors:** Júlia Vallvé-Juanico, Ashley F. George, Sushmita Sen, Reuben Thomas, Min-Gyoung Shin, Divyashree Kushnoor, Joshua J. Vásquez, Kim Chi Vo, Juan C. Irwin, Nadia R. Roan, Alexis J. Combes, Linda C. Giudice

**Affiliations:** Center for Reproductive Sciences, Department of Obstetrics, Gynecology and Reproductive Sciences, University of California San Francisco, San Francisco, CA, USA; Bioliquid Innovative Genetics S.L., Barcelona, Spain; Gladstone Institutes, San Francisco, CA, USA; Department of Urology, University of California San Francisco, San Francisco, CA, USA; San Jose State University, San Jose, CA, USA; Bioinformatic Core Gladstone Institutes, San Francisco, CA, USA; UCSF CoLabs, University of California of San Francisco, San Francisco, CA, USA; ImmunoX initiative, University of California of San Francisco, San Francisco, CA, USA; Department of Medicine, University of California of San Francisco, San Francisco, CA, USA; Department of Pathology, University of California of San Francisco, San Francisco, CA, USA

**Keywords:** endometriosis, macrophages, monocytes, mononuclear phagocytes, SIRPα, innate immune, CyTOF, biomarker, endometrium

## Abstract

**Background:** Endometriosis is a chronic, estrogen-dependent disorder where inflammation contributes to disease-associated symptoms of pelvic pain and infertility. Immune dysfunction includes insufficient immune lesion clearance, a pro-inflammatory endometrial environment, and systemic inflammation. Comprehensive understanding of endometriosis immune pathophysiology in different hormonal milieu and disease severity has been hampered by limited direct characterization of immune populations in endometrium, blood, and lesions. Simultaneous deep phenotyping at single cell resolution of complex tissues has transformed our understanding of the immune system and its role in many diseases. Herein, we report mass cytometry and high dimensional analyses to study immune cell phenotypes, abundance, activation states, and functions in endometrium and blood of women with and without endometriosis in different cycle phases and disease stages.

**Methods:** A case-control study was designed. Endometrial biopsies and blood (n=60 total) were obtained from women with (n=20, n=17, respectively) and without (n=14, n=9) endometriosis in the proliferative and secretory cycle phases of the menstrual cycle. Two mass cytometry panels were designed; one broad panel and one specific for mononuclear phagocytic cells (MPC), and all samples were multiplexed to characterize both endometrium and blood immune composition at unprecedented resolution. We combined supervised and unsupervised analyses to finely define the immune cell subsets with an emphasis on MPC. Then, association between cell types, protein expression, disease status, and cycle phase were performed.

**Results:** The broad panel highlighted a significant modification of MPC in endometriosis; thus, they were studied in detail with an MPC-focused panel. Endometrial CD91^+^ macrophages overexpressed SIRPα (phagocytosis inhibitor) and CD64 (associated with inflammation) in endometriosis, and they were more abundant in mild versus severe disease. In blood, classical and intermediate monocytes were less abundant in endometriosis, whereas plasmacytoid dendritic cells and non-classical monocytes were more abundant. Non-classical monocytes were higher in severe versus mild disease.

**Conclusions:** A greater inflammatory phenotype and decreased phagocytic capacity of endometrial macrophages in endometriosis are consistent with defective clearance of endometrial cells shed during menses and in tissue homeostasis, with implications in endometriosis pathogenesis and pathophysiology. Different proportions of monocytes and plasmacytoid dendritic cells in blood from endometriosis suggest systemically aberrant functionality of the myeloid system opening new venues for the study of biomarkers and therapies for endometriosis.

## Background

Endometriosis is a common, chronic, estrogen-dependent, inflammatory disorder affecting approximately 10% of reproductive age women, 60% with chronic pelvic pain, and 50% with infertility [1]. It is characterized by endometrial-like tissue outside the uterine cavity, arriving in the pelvis by retrograde menstruation and at distant sites by hematogenous and/or lymphatic spread [2]. Endometrial tissue at sites ectopic to the uterus results in an intense inflammatory response, neo-neuroangiogenesis, fibrosis, and scarring [1]. About 97% of women have retrograde menstruation, although most do not develop endometriosis, and while the risk of developing disease is approximately 50% genetic and 50% environmental [1], predicting who will develop symptoms remains uncertain. There are no biomarkers for endometriosis and the gold-standard diagnosis is by laparoscopy. Current therapies include surgical removal of lesions or hormonal ovarian suppression, but disease recurrence and medication side-effects limit long-term symptom relief [1]. Establishment of disease has been attributed to a profound but incompletely understood dysfunction of both innate and adaptive immune responses. The aberrant immune milieu is associated with inefficient clearance of ectopic lesions, pelvic and systemic inflammation, and abnormalities in the endometrium, including alterations in the landscape and function of local immune cells [3, 4]. Understanding endometriosis-related immune dysfunction may be important in defining clinically meaningful disease phenotypes and support the development of novel diagnostic and therapeutic strategies.

Endometriosis has been called “the disease of the macrophage”, based on studies revealing a central role for macrophages in disease establishment, angiogenesis, nerve fiber development and involvement in pain perception [5]. Recently, a mouse model of the disease demonstrated that macrophages in lesions derive from eutopic endometrium (lining the uterus), circulating monocytes, and large peritoneal macrophages [6]. Moreover, it demonstrated that endometrial macrophages are “pro-endometriosis”, as their depletion resulted in smaller endometriotic lesions, while monocyte-derived peritoneal macrophages are “anti-endometriosis”, protecting from lesion establishment [6]. These data confirm key roles for endometrial macrophages and a possible defect in myeloid function in this disorder. Recently, we found, by RNA-Seq, that isolated endometrial macrophages from women with endometriosis have signatures of enhanced inflammation, supporting a role for these cells in the pathogenesis and pathophysiology of this disorder [7]. Molecular and cellular characterization of the human myeloid system in peripheral blood and endometrium of women with endometriosis has been limited, in part, by established methods to isolate and study these cells. Technologic advances in immunology research, including mass cytometry (i.e., *Cytometry by Time-Of-Flight* (CyTOF)), enable deep phenotyping of immune and other cell types. Mass cytometry enables simultaneous measurement of more than 40 proteins at single cell resolution in a complex tissue. Herein, we have leveraged CyTOF to study human endometrium and peripheral blood immune cells (PBMCs) of those with endometriosis (cases) and without disease (controls) and in different menstrual cycle phases (estrogen-dominant proliferative phase and progesterone-dominant secretory phase). Two complementary CyTOF panels consisting of 42 and 40 phenotypic and functional markers were designed. The first, a broad immune phenotyping panel, enabled resolution and comparison of endometrial immune populations. The second panel focused exclusively on phenotyping mononuclear phagocytic cells (MPC) in both endometrium and blood. The data reveal enrichment and activation of distinct populations in cases (endometriosis) and controls, menstrual cycle phases, and endometriosis disease stages and offer candidates for diagnostic and therapeutic target development.

## Methods

### Sample collection and processing

A total of 34 fresh endometrial tissues (Control = 14, Endometriosis = 20) and 26 blood samples (Control = 9, Endometriosis = 17) were collected from women with and without endometriosis undergoing surgery (total n = 60). Clinical features of the patients (cycle phase, diagnosis, and stage of disease) are listed in Supplemental Table 1. Diagnosis was made by the physician and pathologists at the University of California, San Francisco (UCSF), and stage of disease was determined following the rASRM classification system (stages I and II were determined as mild and III and IV as severe stages) [8]. Women without visualized endometriosis at the time of surgery or without a history of endometriosis were defined as controls. Women with any type of cancer and/or endometrial hyperplasia were also excluded. Some patients exhibited non-malignant gynecologic disorders such as leiomyomas or uterine polyps. Cycle phase was determined by following Noyes et al. system of classification of endometrial histology [9]. Clinical features of the patients were collected only by authorized personnel by using the REDCap Database, after informed written consent. Only patients of reproductive age (18-49 years old) were included in the study. In addition, patients presenting with immune-related comorbidities, such as systemic lupus erythematosus (SLE), endometritis, or other immune disorders, were excluded, as well as those positive for HIV, HVB and HVC. Moreover, patients were not exposed to hormone therapies for at least 3 months prior to biospecimen collection. Finally, women under any treatment containing iodine were also excluded from the study, as this element interferes with the CyTOF instrument. All samples were obtained between 2019 and 2021 under the auspices of the UCSF Institutional Review Board Procotol #: IRB#10-03964, using the WERF EPHect standardized protocols for tissue collection, and processing, and clinical annotation [10]. All patient data were de-identified and followed HIPAA and the Convention of the Declaration of Helsinki.

#### Endometrial samples

Endometrium was obtained either by endometrial biopsy using a Pipelle catheter (CooperSurgical, Trumbull, CT, USA) or from hysterectomy specimens. Tissues were placed into transport medium and processed within 5 hours following collection where they were first washed with serum containing media (SCM) and then digested mechanically and enzymatically using a mix of collagenase IV and hyaluronidase, as previously described [7]. After one hour of digestion at 37°C under rotation, for live/dead discrimination, samples were processed for incorporation of cisplatin, and fixed for further usage. Briefly, the single cell suspension was washed with FACS/EDTA buffer (PBS supplemented with 2% FBS and 2mM EDTA). Cells were counted and an appropriate amount of cisplatin (Fluidigm, South San Francisco, CA, USA) (25mM per 1-6 million cells) was added to the suspension (4ml PBS/EDTA per 1-6 million cells) for exactly 60 seconds at room temperature (RT). Then the cells were quenched with CyFACS (metal contaminant-free PBS (Rockland, Pottstown, PA, USA) supplemented with 0.1% BSA and 0.1% sodium azide). Finally, cells were fixed with 1.6% formaldehyde for 10 minutes, washed three times in CyFACS, and stored at −80°C until further use.

#### Peripheral blood mononuclear cells (PBMCs)

Blood was collected in collection tubes containing anticoagulant acid citrate dextrose (ADC) Solution B (Fisher Scientific, Hampton, NH, USA). Ficoll (Stemcell Technologies, Vancouver, Canada) was slowly added to the bottom of the blood in a falcon tube at a ratio of 2:1. The samples were then centrifuged at 2000 rpm for 30 minutes at RT. After centrifugation, the supernatant was removed and the PBMCs layer was carefully collected. PBMCs were washed twice with FACS buffer, and the number of cells was then counted. The same protocol above was used to incorporate cisplatin, fix, and store the cells.

### Panel designs

First, we designed a CyTOF broad panel to identify the important cell types in endometrial tissue of women with endometriosis compared to controls. This panel consisted of 42 markers and is shown in Supplementary **Table 2.** Then, we designed a 40-parameter CyTOF panel that includes mostly myeloid surface markers as well as functional markers, including efferocytosis and phagocytosis, activation, and inhibition markers **(Supplemental Table 2).** For the broad panel, all 42 antibodies were conjugated in house. For the myeloid panel, sixteen of the 40 antibodies required in-house conjugation to their corresponding metal isotope. Metals were conjugated according to the manufacturer’s instructions (Fluidigm, South San Francisco, CA, USA). Briefly, this process comprises loading the metal to a polymer (incubation of 1h at RT). The unconjugated antibody is transferred into a 50kDA Amicon Ultra 500 V-bottom filter (Fisher Scientific, Hampton, NH, USA) and reduced at 37°C with 1:125 dilution of Tris (2-carboxyethyl) phosphine hydrochloride (TCEP) (ThermoFisher, Waltham, MA, USA) for 30 minutes. Then, the column is washed twice with buffer C (Fluidigm, South San Francisco, CA, USA) and the metal-loaded polymer is suspended in 200μl of C-buffer in the 3kDA Amicon Ultra 500ml V-bottom filter (Fisher Scientific, Hampton, NH, USA). The suspension is then transferred to the 50kDa filter containing the antibody and incubated for 1.5h at 37°C. After the incubation time, antibodies are washed three times with W-buffer (Fluidigm, South San Francisco, CA, USA) and quantified for protein content by Nanodrop. Once the concentration was determined, the antibodies were resuspended with Antibody Stabilizer (Boca Scientific, Dedham, MA, USA) at a concentration of 0.2 mg/ml and stored at 4°C. The rest of the antibodies were commercially available (Fluidigm, South San Francisco, CA, USA). Optimal concentrations of all antibodies were performed by different rounds of titrations.

### Barcoding and Cell Staining with Metal Antibodies

The staining protocol was optimized to use each antibody in aliquots of 6 million cells as previously described [11]. Samples were thawed and washed with FACS buffer. Then, cells were counted and since some samples had fewer than 6 million cells, they were barcoded before the staining with the antibodies, following the manufacturer’s instructions (Fluidigm, South San Francisco, CA, USA). Briefly, each sample was incubated with 10μl of each barcode and perm buffer (Fluidigm, South San Francisco, CA, USA) for 30 minutes at RT, and then samples were combined and split into different tubes for the first day of staining. For the staining, samples were blocked using rat, mouse, and human serum for 15 minutes on ice. They were then washed and stained with the primary cocktail of antibodies for 45 minutes at 4°C. After this incubation time, cells were washed and fixed using 2% paraformaldehyde (PFA) diluted in CyPBS. Cells were incubated overnight at 4°C, and the next day were washed with perm buffer (Fluidigm, South San Francisco, CA, USA), washed with CyPBS, and blocked with rat and mouse serum for 15 minutes on ice. They were then washed, and the intracellular staining was performed. Cells were resuspended with the intracellular cocktail of antibodies for 45 minutes on ice and were washed and incubated for 20 minutes at RT with Ir-intercalator (Biolegend CNS, San Diego, CA, USA), prepared at a dilution of 1:500 in 2% fresh PFA. After the incubation time, cells were washed and kept at 4°C overnight. Finally, on the third day, cells were washed with cell staining media (CSM, Fluidigm, South San Francisco, CA, USA), then with water, and then with cell acquisition solution (CAS, Fluidigm, South San Francisco, CA, USA) at RT. Subsequently, cells were counted, resuspended in 1x EQ™ calibration beads (Fluidigm, South San Francisco, CA, USA) and CAS and samples were run in the CyTOF^®^2 instrument (Fluidigm, South San Francisco, CA, USA).

### Data processing

The fcs. files obtained from the instrument were concatenated, normalized to EQ™ calibration beads and de-barcoded using CyTOF software (Fluidigm, South San Francisco, CA, USA). This study is comprised of eight different sample groups: control (Ctrl) and endometriosis (Endo) eutopic endometrium (EM) in the proliferative (PE) and secretory (SE) phases (Ctrl_EM_PE, Ctrl_EM_SE, Endo_EM_PE, Endo_EM_SE) and control and blood (PBMCs) in the proliferative and secretory phases (Ctrl_PBMC_PE, Ctrl_PBMC_SE, Endo_PBMC_PE, Endo_PBMC_SE). Normalized data from the broad panel were imported to FlowJo (BD, Franklin Lakes, NJ, USA) to perform manual gating and we performed unsupervised analysis of the manually gated CD45^+^ cells. We also performed manual gating of the different populations obtained from the focused panel and 8 populations were obtained and are shown in **Supplemental Figure 1.** Then, unsupervised analysis of the manually-gated myeloid cells of interest (including macrophages, monocytes, dendritic cells, and plasmacytoid dendritic cells) was also performed. Finally, statistical analyses for specific markers and populations of the focused panel were performed by manual gating to validate the results from the unsupervised analysis. The datasets generated and analyzed during the current study are available in the Dryad repository (doi:10.7272/Q6Q52MVQ).

### Data and statistical analysis

#### Samples included in the study

The broad panel included a total of 17 endometrial samples (4 controls in the proliferative phase, 2 controls in the secretory phase, 6 from endometriosis patients (cases) in the proliferative phase and 5 cases in the secretory phase). For the focused panel, the endometrial data included 13 control samples (9 in the PE and 4 in SE phases, respectively) and 18 samples from women with endometriosis, which correspond to 13 in the PE phase (8 mild and 5 severe stages, respectively) and 5 in the SE phase (all mild stage of disease). In the case of the PBMCs, data from the focused panel included 9 control samples (6 in the PE and 3 in the SE phase) and 17 disease samples, corresponding to 13 in the PE phase (8 mild stage and 5 severe stage) and 4 in SE phase (3 mild and one severe). Note that some samples were used for both panels, totally 60 in the study (34 endometrial tissues (n=20 control and n=14 endometriosis cases) and 26 blood samples (n=17 controls and n=9 endometriosis cases).

#### Downsampling cells

The broad panel included a total of 17 endometrial samples. Endometrial samples with large numbers of cells were downsized by random cell selection to a number of cells (169,599) per sample compatible with the system’s memory limits and computational efficiency. The Seurat R package for single-cell analysis [12] was used to identify clusters of cells. After combining samples, a total of 2,223,274 endometrial cells were subjected to Seurat clustering, using the levels of the 42 CyTOF markers as expression value to create Seurat objects. CyTOF clusters were identified using a shared nearest neighbor (SNN) graph [13]. Similarly, in the focused panel, 50,000 cells per sample from PBMCs were down sampled from each subject group to enable processing within memory limits and computational efficiency. On the other hand, endometrial samples in the focused panel were not down sampled as they displayed an average of 15,000 mononuclear phagocytic cells per sample after elimination of the lineage positive (CD3+, CD56+, CD66b+). Samples with less than 1,000 CD45+ cells were discarded from the study and are not shown here. Due to the different origin and properties of the two tissues, two independent Seurat objects were created. A total of 355,240 endometrial myeloid phagocytic cells and 890,602 myeloid phagocytic cells from blood were subjected to Seurat clustering. The expression level of the markers from the panel were given as expression value to create Seurat objects.

As only myeloid populations were gated and analyzed, markers for other cell types were excluded from the analysis (CD45, CD3, CD56, CD66b). In addition, the expression levels of the antibodies MerTK and Erα were negative, indicating that the staining did not perform well, and these two markers were also excluded from the analyses, resulting in the inclusion of 34 markers from the panel to the final analysis. The downsampling of cells should not have resulted in any bias because cells from all subjects in the study were included in the analyses and subjected to an unbiased downsampling. Moreover, cells only from subjects with numbers of cells exceeding a chosen threshold (depending on the sample type as explained above) were downsampled.

#### Visualization using UMAPs colored by biological variables and technical variables

Biological variables (disease, menstrual cycle phase) and technical variables (batch/run) were visualized in different colors using the DimPlot function in the Seurat package. Because two sampling methods were used to collect the endometrial samples (hysterectomy and biopsy), we also generated and compared UMAP coordinates of the cells between the two collection methods. We concluded that they were comparable, and thus all endometrial samples were used **(Supplemental Figure 2).**

#### Batch correction procedure using Harmony

After visualizing batch effects in the broad panel using DimPlot, the first and remaining group of the three batches were treated as two batch groups and the cells were combined using the harmony batch correction function RunHarmony [14]. Similarly, samples run in batch/run 5 for the focused panel (both endometrial and PBMCs) were systematically different from the samples run in batches 1-4 **(Supplemental Figure 3).** Therefore, samples under run 5 and the rest of the samples were treated as two separate batches that were subjected to correction using the RunHarmony function to proceed with further analysis.

#### Clustering of cells at different resolutions

Clustering of the cells was performed using the FindClusters function (implementing the “original Louvain” algorithm) in the Seurat package at resolutions 0.01, 0.1, 0.2, 0.4, 0.8, 1 and 1.5. At each of these resolutions, the markers for each cluster were determined using the FindMarkers function in Seurat where the processing batch for each sample was encoded as a latent variable. The average expression for each marker was determined using the AverageExpression function in the Seurat. After looking at the cisplatin levels (dead cells), some clusters were removed, as they presented high levels of dead cells; also, very small clusters that contained less than 100 cells were excluded. In the broad panel, the resolution parameter was chosen to be 0.2 for endometrium. In the focused panel, the resolution parameter was chosen to be 0.4 and 0.2 for endometrium and PBMCs, respectively.

#### Between cluster association with disease and menstrual cycle state

A generalized linear mixed model (GLMM) (implemented in the lme4 [14] package in R with family argument set to the binomial probability distribution) was used to estimate the association between cluster membership, disease, and menstrual cycle phase. The model consisted of cluster membership as a response variable and five explanatory variables: subject as a random effect variable, variables encoding disease, menstrual cycle phase, their interaction, and a batch variable were included as fixed variables.

#### Between cluster association with disease stage

GLMM was also used to explore the association between cluster membership and disease stage of cells. Disease samples from patients with mild or severe stage were selected for disease stage association analysis. The model explanatory variables included subject as a random effect, disease stage, menstrual cycle, and the processing batch (for analyses involving disease samples processed across multiple batches) as fixed variables. Only proliferative phase samples were studied due to limited secretory phase samples from patients with severe disease.

#### Within cluster association with disease and menstrual cycle

To assess the association (per cluster) between CyTOF marker expression quantile levels and disease, menstrual cycle phase and their interaction, a linear model was fit. The tested marker expression quantile values of 25%, 50%, and 75% for each subject were estimated across all the subjects’ cells. In addition to variables that capture disease status, menstrual cycle phase, and their interaction, a variable capturing the processing batch was also included as explanatory variables in the linear model.

#### Within cluster association with disease stage

Similarly, a linear model was used to explore the association between CyTOF marker cell expression quantile levels and disease stage. The variables included in these models are described in the section on “Between cluster association with disease stage". All p-values were corrected using the false discovery rate (FDR), and a threshold of 0.05 was used to determine significance for the distribution of cells in each cluster and 0.1 for marker expression. GLMM was performed using R package lme4 [15], and linear regression was performed using the lm function in R. Only proliferative phase samples were studied.

#### Validation of the unsupervised analysis by manual gating

To validate the results, we manually gated the populations of interest in FlowJo and studied their abundance and marker expression. To find differences in expression between the manually gated populations, we extracted the mean signal intensity (MSI) of each marker, and its expression was compared between the different groups by t-test (p ≤ 0.05) and multiple correction with Benjamini-Hochberg (BH) with a false discovery rate (FDR) lower than 0.05.

## Results

### Broad immune phenotyping identifies differences in abundance and variation throughout the cycle in endometrial immune populations in women with and without endometriosis

Clustering analysis (which include all samples analyzed in the study; controls and cases with endometriosis in both proliferative and secretory phases of the cycle) revealed 11 distinct immune clusters and one cluster of putative endothelial cells, the latter of which has been observed to express CD45 at low levels in some tissues [16]. The 11 immune cell clusters correspond to macrophages, natural killer cells (NK), neutrophils, and subsets of CD4^+^ T cells including Temra (effector memory T cells that re-express CD45RA after antigenic stimulation), CD8^+^ T cells, CD16^+^ NK, CD69+ NK, B cells, γδ T cells, and class I conventional dendritic cells (cDC1) **(Figure 1).** Briefly, T cells were identified by the high expression of CD3. Their subsets were identified by the high expression of CD4 (T cells CD4+) and CD8 (T cells CD8+), CD45RA (Temra), and γδTCR (γδ T cells). B cells were identified by the expression of CD19. Cells expressing high levels of CD15 and CD126 were classified as neutrophils. Macrophages were identified by the high expression of HLA-DR, CD11c, CD36, and CD14, and they were negative for CD3, CD56, and CD19, among others. We found three clusters of natural killer cells (NK). We identified them as they are all are negative for the lineage markers CD3, CD16, and CD14 but express different levels of CD56. In addition, the close proximity of clusters 1, 7 and 8 on the UMAP indicates similarities among the three clusters. So, to differentiate among the three NK cells subsets, we named them with the expression of the most distinguishing markers based on the literature. First, Cluster 1 (NK) is CD56+ and CD16−, which corresponds to infiltrating natural killers (from blood); cluster 7 (NK CD16+) corresponds to endometrial natural killers (CD56 dim and CD16 bright, markers consistent with activation; and third, those that highly express CD69, a marker previously described in tissue-resident NK cells [17]. Endothelial cells were defined by the high-level expression of BDCA3. Cluster 11 was annotated and defined as cDC1 due to its expression of BDCA3, CD36, CD11c, and HLA-DR and being negative for CD3, CD4, CD14, and BDCA1, which are gold standard markers for cDC1 in humans. Nonetheless, the high BDCA3 expression in endothelial cells and the normalization across the column in the heatmap obscure the expression of BDCA3 in cluster 11 **(Figure 2b).** The proportion of each population in all conditions (control and endometriosis in both phases of the cycle) is shown in **Supplemental Figure 4.** Populations displaying significant differences in abundance between controls and cases across the menstrual cycle are presented in **Figure 2. A.** The distribution of all populations in both controls and cases is shown in **Supplemental Figure 5.** In controls, the abundance of cDC1 significantly increased in the secretory phase of the cycle as did macrophages albeit insignificantly, while in women with endometriosis, both populations significantly decreased **(Figure 2. A).** In addition, while B cells tended to decrease in the secretory phase in controls, this population significantly more abundant in samples acquired during the secretory phase from cases **(Figure 2. A).** Furthermore, we found statistically significant differences between abundance of specific populations in cases and controls. We observed increased abundance of macrophages and neutrophils in the proliferative phase and decreased CD69+ NK, B cells, and γδ Tcells in the proliferative phase in cases vs controls **(Figure 2. B).**

**Figure 1.**
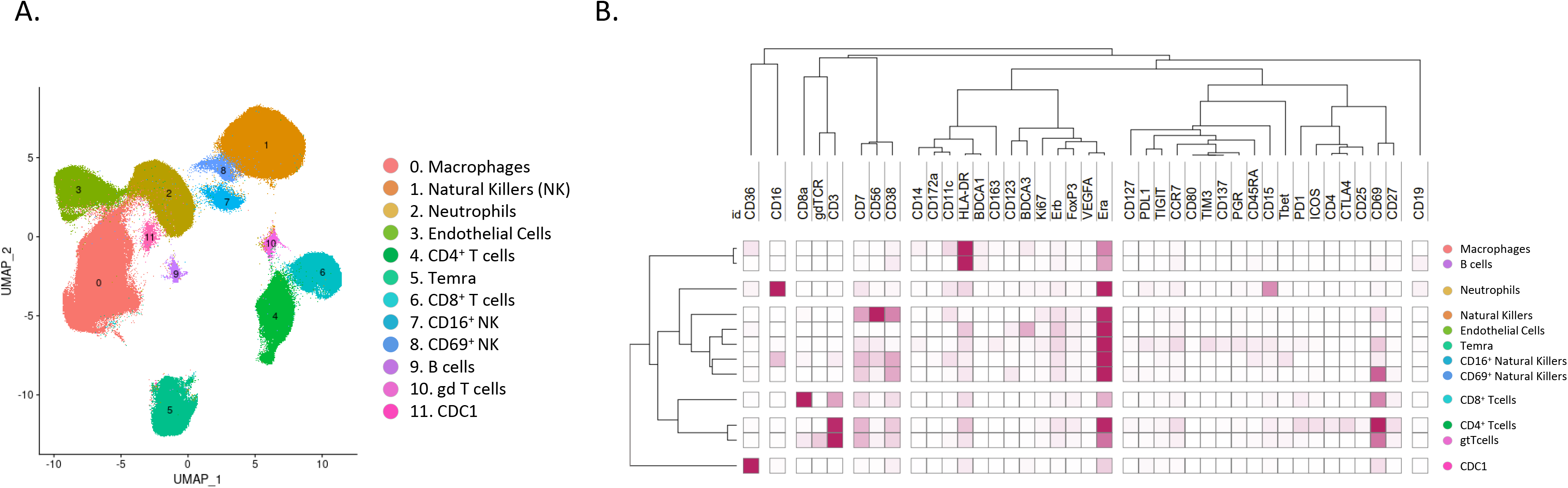
Unsupervised clustering using broad immune panel identified 12 unique cell population in endometrium. **A)** UMAP showing the 12 identified clusters. **B)** Heatmap with the marker’s level expression in each of the populations identified from the clustering. n=17 (4 controls PE, 2 controls SE, 6 endometriosis PE, and 5 endometriosis SE). PE: proliferative, SE: secretory.

**Figure 2.**
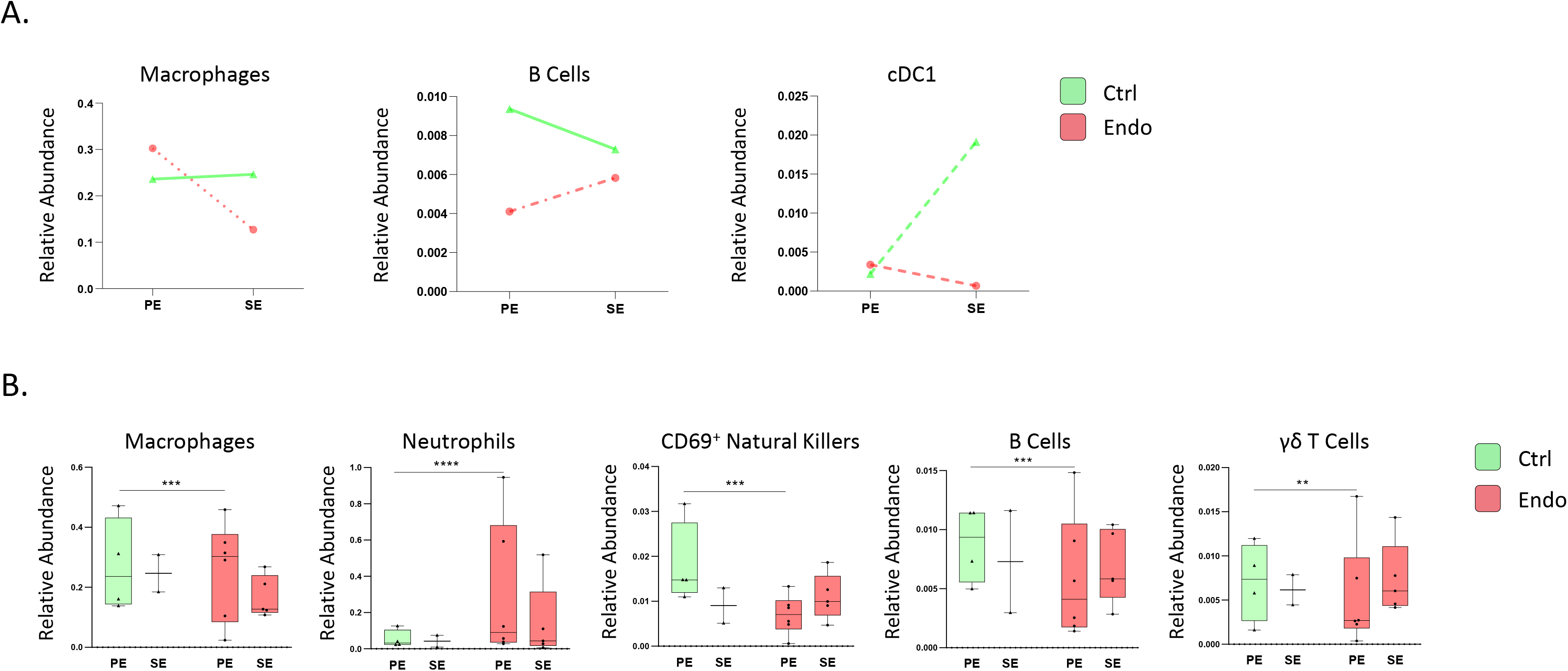
Broad panel association analysis in endometrium. **A)** The figure shows only the populations that vary significantly different throughout the cycle between controls and endometriosis patients. pValues: dotted and dashed lines, pVal ≤ 0.05; dashed lines, pVal ≤ 0.0005; dotted lines, pVal ≤ 0.00005; straight lines: no significant. **B)** Significant differences in abundance between cases and control immune populations. All analyses were performed in both phases of the menstrual cycle (PE: proliferative, SE: secretory). p Values: *, pVal ≤ 0.05; **, pVal ≤ 0.005; ***, pVal ≤ 0.0005; **** pVal ≤ 0.00005. n=17 (4 controls PE, 2 controls SE, 6 endometriosis PE, and 5 endometriosis SE). PE: proliferative, SE: secretory.

Increased abundance of subsets of macrophages and neutrophils in endometrium of women with endometriosis in the proliferative phase **(Figure 2. B)** indicates more inflammation in tissues from women with disease compared to controls. This observation suggesting that myeloid populations may be involved in disease prompted the development of a mononuclear phagocyte-focused panel to study these populations in more detail.

### Mass cytometry reveals distinct subsets of mononuclear phagocyte populations in endometrium

We then combined phenotypic and functional markers of the mononuclear phagocyte populations including activation/inhibition markers, phagocytosis markers, and efferocytosis markers. By using manual gating, we first excluded T Cells, NK, and granulocytes, and then performed the unsupervised analysis on the remaining cells. The unsupervised clustering yielded 13 unique cell populations **(Figure 3),** including one which was characterized by an absence of expression of any mononuclear phagocyte markers and was therefore excluded for further analysis, as it may be a potential B cells cluster. In this manner, we identified 12 clusters in the endometrial tissue (analysis made with all samples; cases and controls in both proliferative and secretory phases). We found three clusters corresponding to classical monocytes (CD44^+^ classical monocytes, HLA-DR^+^ classical monocytes, and HLA-DPB1^+^ classical monocytes) defined by a high expression of CD14 and CD36, and low expression of CD16; three macrophage clusters (CD206^+^ macrophages, CD91^+^ macrophages, and ALXR^+^ macrophages) defined by the expression of CD11c, CD64, and CD206; two clusters of non-classical monocytes (CX3CR1^+^ non-classical monocytes and non-classical monocytes) defined by their CD16 expression and low expression of CD14. In the case of the dendritic cells’ classification, the focused panel did not contain a great number of specific markers for subsets of dendritic cells, such as CD141 or CD1c, which made more difficult their identification. However, it included CD1a, CD11c, CD123, CD14, and CD16 which allowed their broad identification. Consequently, we used a hybrid strategy to identify DCs subsets. We first isolated plasmacytoid dendritic cells (pDCs) using the gold standard classification of lineage negative (CD3, CD56, CD14, CD16−, and CD11c−) and expression of HLA-DR+, CD11c−, CD16−, and CD123+ markers. Regarding classical Dendritic Cells, we had not included in this panel the markers BDCA1 (CD1c) and BDCA3 (CD141), classical markers for cDC2 and cDC1, respectively. Nevertheless, based on the literature, cDCs are CD14dim, and CD11c+ (also shown in Supplemental Figure 1), which is the case for the three clusters of dendritic cells that we have identified named dendritic cells, TGM2+ dendritic cells and CD1a+ dendritic cells. In addition, we added dendritic cell markers in the focused panel that we believed could be involved in endometriosis. It is well known that dendritic cells can express CD1a+ [18]. This marker is believed to be an immature marker for DCs while CD83+ is a marker of mature DCs. As it was previously described that in endometrium of women with endometriosis, immature DCs (CD1a+) were more abundant than CD83+ mature DCs [19], these two markers were included in the focused panel. We identified a cluster (cluster 5) with high expression of CD1a, indicating that these may be immature dendritic cells. On the other hand, TGM2+ is also expressed in DCs [20] and it is believed to be involved in dendritic cell-T cell interaction. However, it is inevitable that the relationship between those subsets and the classical cDC1 and cDC2 will need to be established.-The expression levels of all markers for each population are shown in **Figure 3. B and** in **Table 1.**

**Figure 3.**
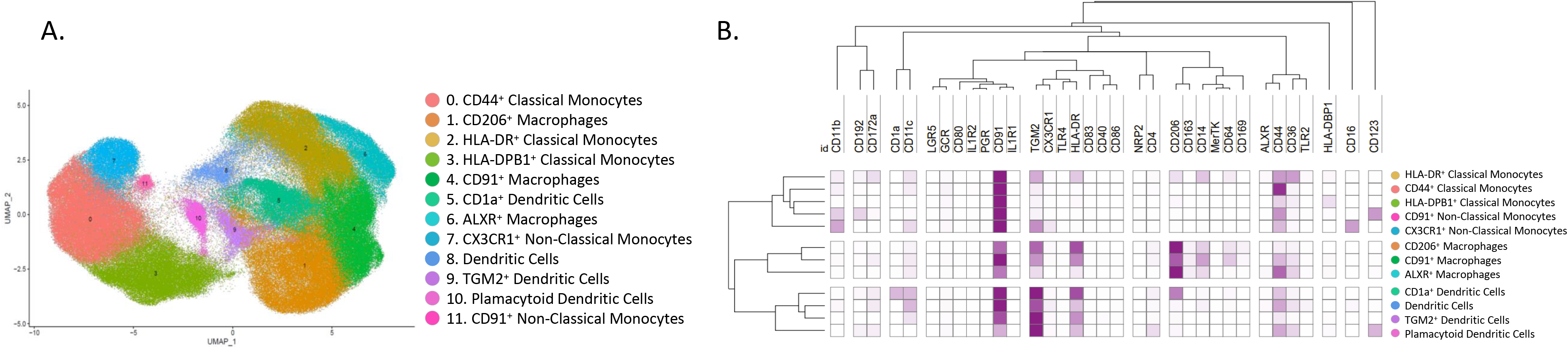
Identified populations from the unsupervised clustering analysis in endometrium in the focused panel. **A)** UMAP showing the 12 identified clusters. **B)** Heatmap with the marker’s relative level expression in each of the populations identified from the clustering. n=13 controls (9 PE and 4 SE) and n=18 endometriosis (13 PE (8 mild and 5 severe stages) and 5 SE (all mild stage of disease)). PE: proliferative, SE: secretory.

**Table 1.**
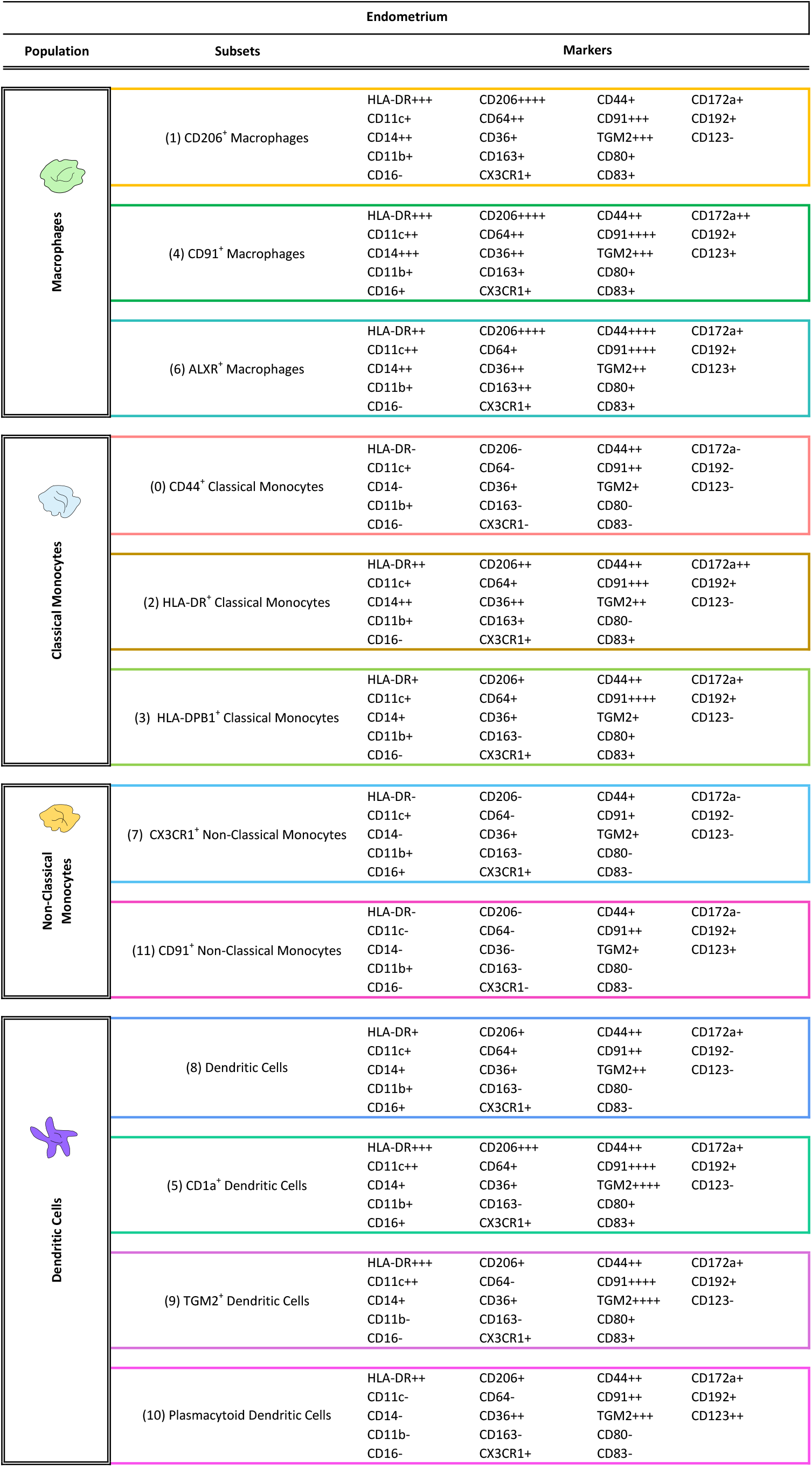
Identified populations from the unsupervised clustering analysis obtained from the focused panel. The numbers in the parenthesis indicate the number of each cluster showed in the UMAP. Levels of expression were determined by the intensity mean of each marker in each cluster; 0-5 (−), 6-100 (+), 101-500 (++), 501-1000 (+++),> 1001 (++++).

### Specific endometrial monocyte and macrophage subsets display distinct dynamics throughout the menstrual cycle in endometriosis patients

In the endometrium of controls, CD44^+^ classical monocytes and ALXR^+^ macrophages decreased in the secretory phase of the cycle, whereas HLA-DR^+^ classical monocytes and CD91^+^ macrophages increased **(Supplemental Figure 6).** Interestingly, the opposite pattern was observed in CD44+ classical monocytes and CD91^+^ macrophages from women with endometriosis **(Figure 4. A).** The distribution of all identified populations with the focused panel in both groups is shown in **Supplemental Figure 6.** These observations indicate that the specific populations have different dynamics throughout the cycle between cases and controls, suggesting different hormonal regulation of these populations in disease.

**Figure 4.**
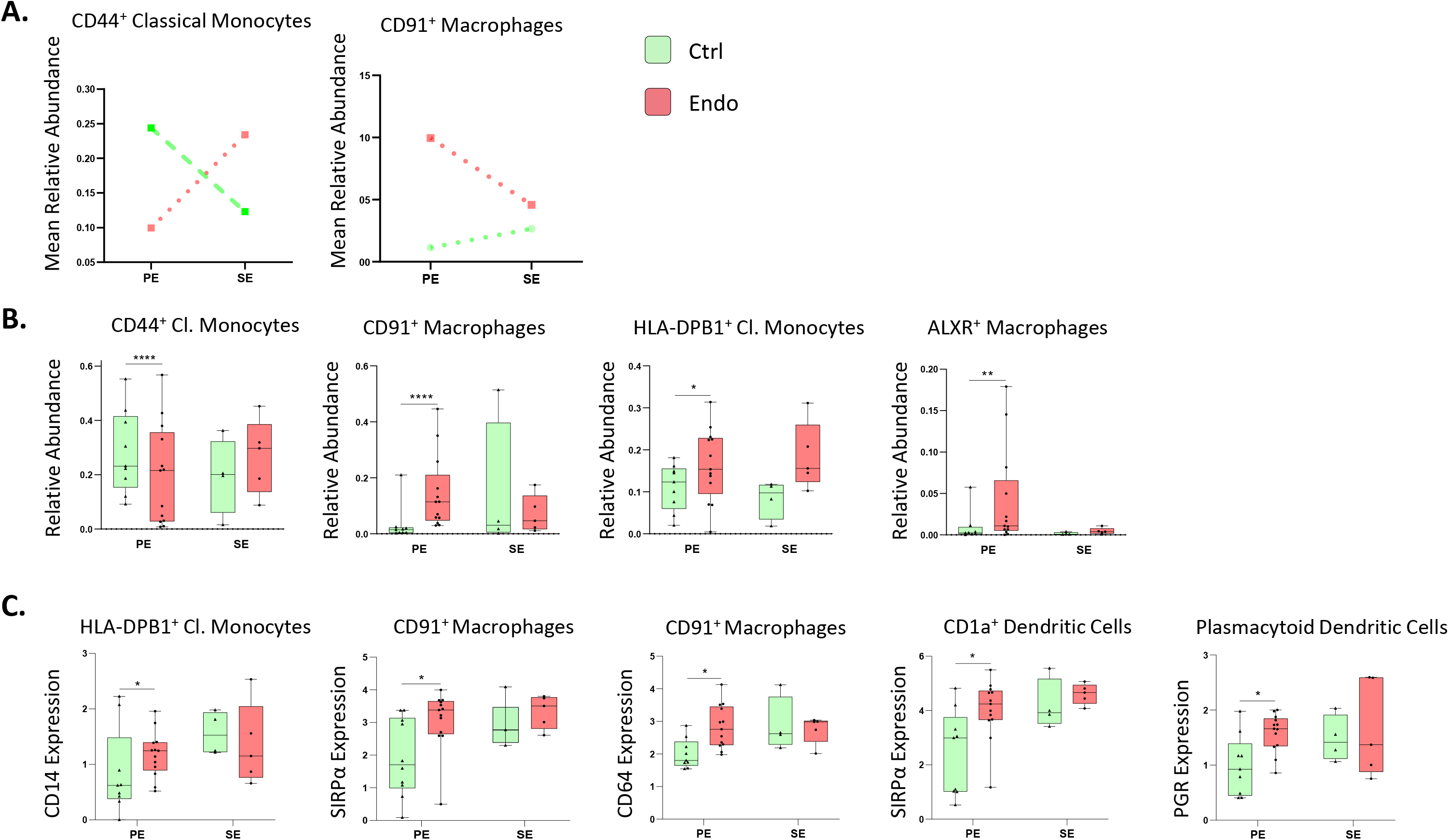
Focused panel association analysis of endometrium in cases and controls and different cycle phases. **A)** Populations with significantly different relative abundance between proliferative and secretory phases in control and endometriosis patients and showing contrary fluctuation between both groups (mean relative abundance is shown). pValues: dashed lines, pVal ≤ 0.0005; dotted lines, pVal ≤ 0.00005. **B)** Significant differences in abundance of immune populations between control and endometriosis patients. pValues: *, pVal ≤ 0.05; **, pVal ≤ 0.005; ***, pVal ≤ 0.0005; **** pVal ≤ 0.00005. **C)** Significant differences in marker expression in immune populations between control and endometriosis patients (pVal ≤ 0.1). n=13 controls (9 PE and 4 SE) and n=18 endometriosis (13 PE (8 mild and 5 severe stages) and 5 SE (all mild stage of disease)). Green: controls; red: cases. All analyses were performed in both phases of the menstrual cycle. PE: proliferative, SE: secretory.

### Abundance and marker expression of specific proliferative phase mononuclear phagocytic subsets differ in endometriosis patients

Association analysis revealed an increased endometrial HLA-DPB1^+^ classical monocytes, CD91^+^ macrophages, and ALXR^+^ macrophages, and decreased CD44^+^ classical monocytes in cases versus controls in the proliferative phase **(Figure 4. B)**. Within some populations, association analysis revealed several markers significantly different between cases and controls **(Figure 4. C)**. For example, CD14 expression on HLA-DPB1^+^ classical monocytes was increased in the proliferative phase in cases versus controls. Interestingly, the anti-phagocytosis marker SIRPα (CD172a) was increased on phagocytic CD91^+^ macrophages and CD1a^+^ dendritic cells in cases versus controls in the proliferative phase, suggesting decreased phagocytic capacity of these populations in endometriosis patients. In addition, an increase in expression of CD64 was found in proliferative phase in CD91^+^ macrophages from cases, indicating higher inflammation. PGR was more highly expressed in pDCs in cases versus controls in the proliferative phase. These results suggest that specific phagocytic cell subsets may have altered functions in endometrium of women with endometriosis. The expression of the significant markers between cases and controls in all cell types during the proliferative phase is shown in **Supplemental Table 3.**

### Endometrial CD91^+^ macrophages and CD1a^+^ dendritic cells display phenotypes suggesting reduced phagocytic capacity and CD91^+^ macrophages display an inflammatory bias in cases versus controls

Unsupervised analyses showed an increase of SIRPα in CD91^+^ macrophages and in CD1a^+^ dendritic cells, as well as increased levels of CD64 in CD91^+^ macrophages. By differential expression analyses on manually-gated mononuclear phagocytic subsets, we found that, in the proliferative phase, a greater frequency of CD91^+^ macrophages in endometrium of women with endometriosis express SIRPα compared to controls **(Figure 5. A)** and they have higher expression levels of CD64 in the proliferative phase of the menstrual cycle **(Figure 5. B),** corroborating the results observed with the unsupervised analysis. We also validated by manual gating that SIRPα was overexpressed by CD1a^+^ dendritic cells in endometrium of cases versus controls in the proliferative phase **(Figure 5. C)**. These results suggest inhibition of the phagocytic capacity of endometrial CD91^+^ macrophages and CD1a^+^ dendritic cells and an inflammatory bias among CD91^+^ macrophages in women with endometriosis.

**Figure 5.**
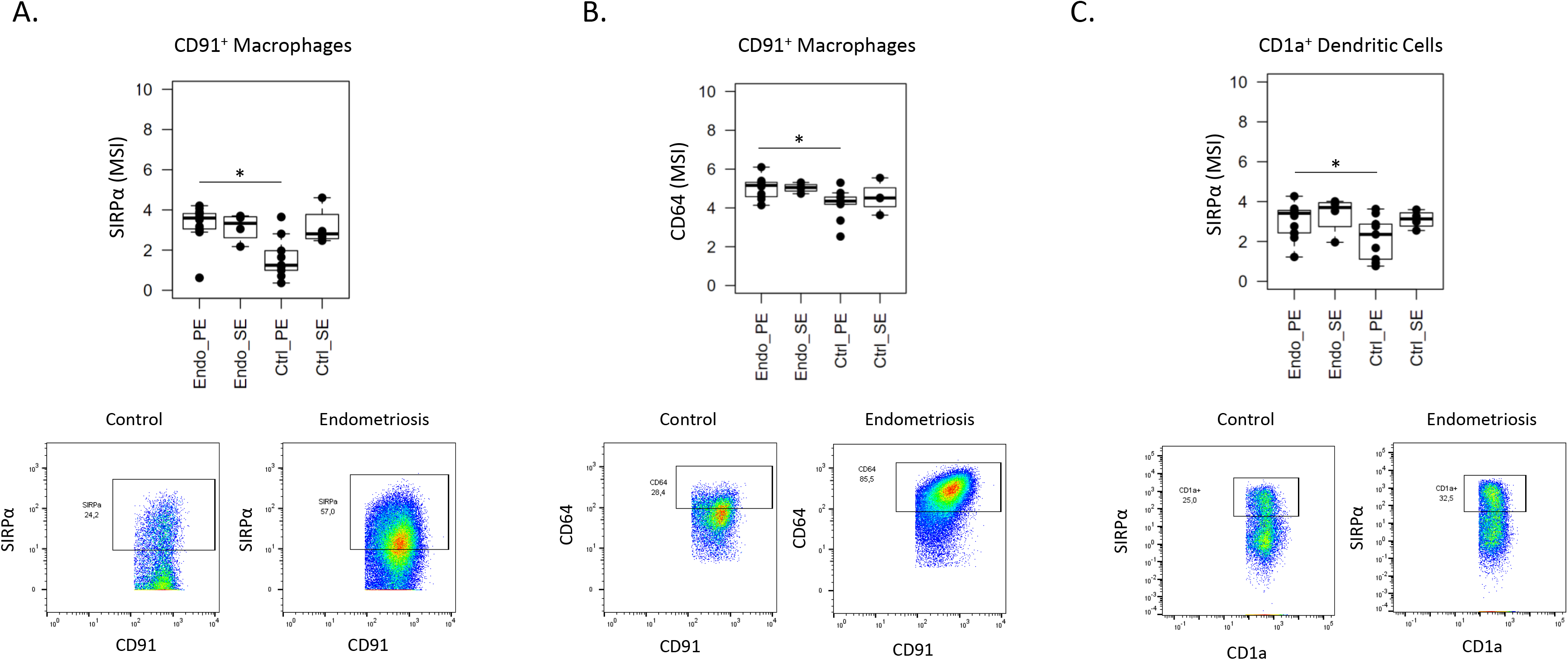
Differentially expressed SIRPα and CD64 in CD91^+^ macrophages and SIRPα in CD1a^+^ dendritic cells from endometrium of women with versus without endometriosis. The figure shows the box plots from the mean signal intensity (MSI) obtained by manual gating using FlowJo^®^ (pValue ≤ 0.05) and the dot plots show the number of cells and intensity of each marker. **A)** SIRPα expression in CD91^+^ macrophages. **B)** CD64 expression in CD91^+^ macrophages. **C)** SIRPα expression in CD1a^+^ dendritic cells. n=9 controls PE and n=13 endometriosis PE. PE: proliferative.

### Distinct endometrial mononuclear phagocyte populations display disease-stage specific abundance in the proliferative phase

In the setting of severe versus mild disease, a higher proportion of endometrial CD44^+^ classical monocytes, CX3CR1^+^ and CD91^+^ non-classical monocytes, and dendritic cells were observed, as well as lower levels of CD91^+^ macrophages **(Figure 6. A).** Notably, and as shown above, CD91^+^ macrophages and CD44^+^ monocytes appear to have different dynamics throughout the cycle in controls and endometriosis patients, as well as different proportions between the two groups, suggesting that they may be key components for the pathophysiology of the disease.

In addition, association analyses of marker expression levels across populations were conducted to identify functional phenotypes and possible differences and roles in the setting of severe versus mild stage disease. This revealed several markers significantly differentially expressed in some endometrial immune populations depending on disease stage **(Figure 6. B).** CD206^+^ macrophages had higher CD80 expression and CD91^+^ macrophages had higher CD64 expression levels in endometrium from women with mild versus severe disease, suggesting higher activation and inflammatory phenotype of these populations in mild disease. Similarly, HLA-DPB1^+^ classical monocytes had higher expression of CD14 and activated classical monocytes (CD44^+^) showed an increase of CD80, CD91, and IL1R2 expression, also indicating higher activation in mild disease. Highly activated classical monocytes (HLA-DR^+^) displayed increased CD83, MerTK, LGR5, CD80, GCR, CD91, IL1R2, CD36, and CD4, showing also higher activation in mild stage. CX3CR1^+^ non-classical monocytes showed an increase in CD91 in patients with mild endometriosis. Dendritic cells had higher expression of CD192 (CCR2), CD80, GCR, CD91, IL1R2 and CD4 and a decrease of ALXR, suggesting that some of these cells are monocyte-derived dendritic cells and that they are more activated in mild than in severe disease. Finally, TGM2^+^ dendritic cells presented a higher expression of IL1R1 in endometrium of women with mild versus severe disease. Taken together, these results indicate that there is more activation in specific immune populations in mild disease when compared to severe disease, suggesting more inflammation endometrium in early stages of endometriosis.

**Figure 6.**
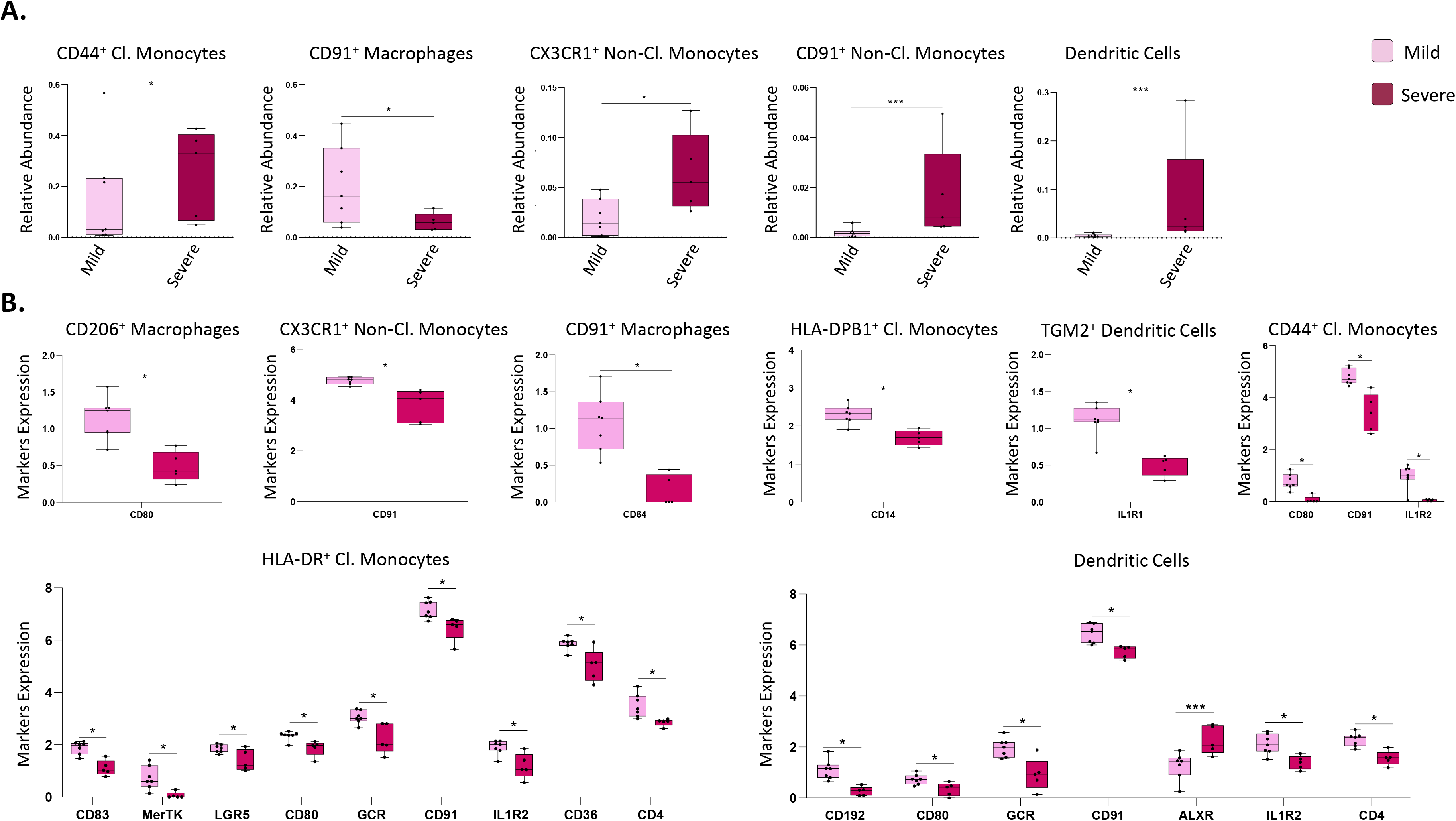
Focused panel association analysis between stages of endometriosis in endometrium. **A)** Differences in abundance of populations between samples from patients with mild and severe disease. pValues: *, pVal ≤ 0.05; **, pVal ≤ 0.005; ***, pVal ≤ 0.0005; **** pVal ≤ 0.00005. **B)** Significant differences in marker expression in specific populations. Both analyses were performed in the proliferative phase of the menstrual cycle. pValues: *, pVal ≤ 0.1; ***, pVal ≤ 0.005. n=13 endometriosis PE (8 mild and 5 severe stages). PE: proliferative.

### Distinct mononuclear phagocyte populations in peripheral blood are associated with endometriosis disease status

As the data in endometrium revealed differences in macrophages, some monocytes, and dendritic cells in women with versus without endometriosis, PBMCs were then analyzed using the focused panel to evaluate possible comparable dysregulation and if these populations could be an indicator of disease. Because only differences in the proliferative phase of the cycle were observed in endometrium, the analysis was restricted to samples from this cycle phase. After the clustering analysis, 11 populations, were identified, corresponding to three clusters of classical monocytes (classical monocytes, activated classical monocytes, and highly activated classical monocytes) defined by their expression of CD14 (high) and CD16 (low) and the expression of activation markers, such as HLA-DR and CD44; three clusters of intermediate monocytes (intermediate monocytes, activated intermediate monocytes, and highly activated intermediate monocytes) also identified by the expression of both CD14 and CD16 and activation markers; three clusters of non-classical monocytes (non-classical monocytes, activated non-classical monocytes, and highly activated non-classical monocytes) defined by their expression of CD14 (low) and CD16 (high) and activation markers; one cluster of dendritic cells, defined by their expression of HLA-DR, CD11c, among others; and one cluster of pDCs, identified by their high expression of CD123. The specific expression of all markers from the panel in each population are shown in **Figure 7** and **Table 2.**

**Figure 7.**
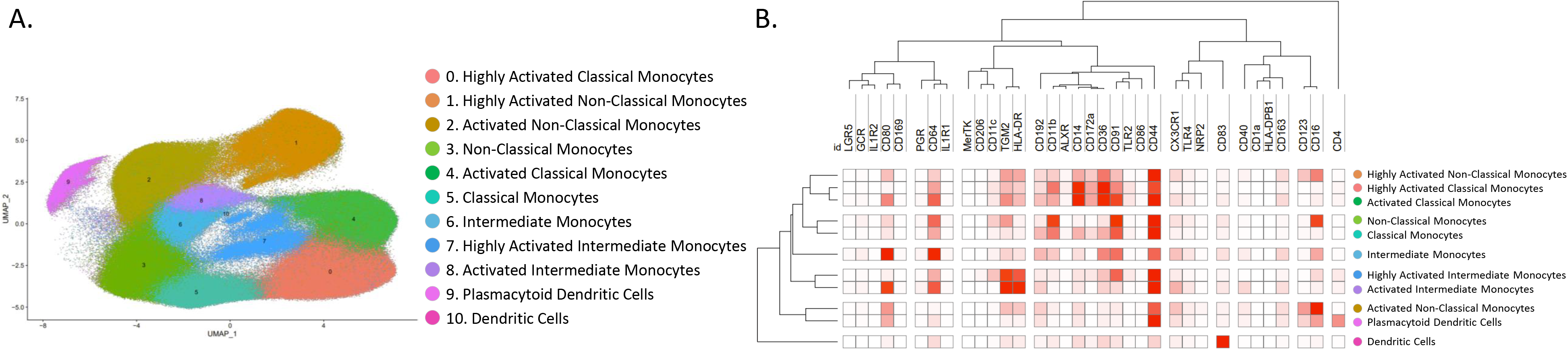
Identified populations from the unsupervised clustering analysis in blood (PBMCs) using the focused panel. **A)** UMAP showing the 11 identified clusters. **B)** Heatmap with the marker level expression in each of the populations identified from the clustering. n=6 controls and n=13 endometriosis PE (8 mild stage and 5 severe stage). PE: proliferative.

**Table 2.**
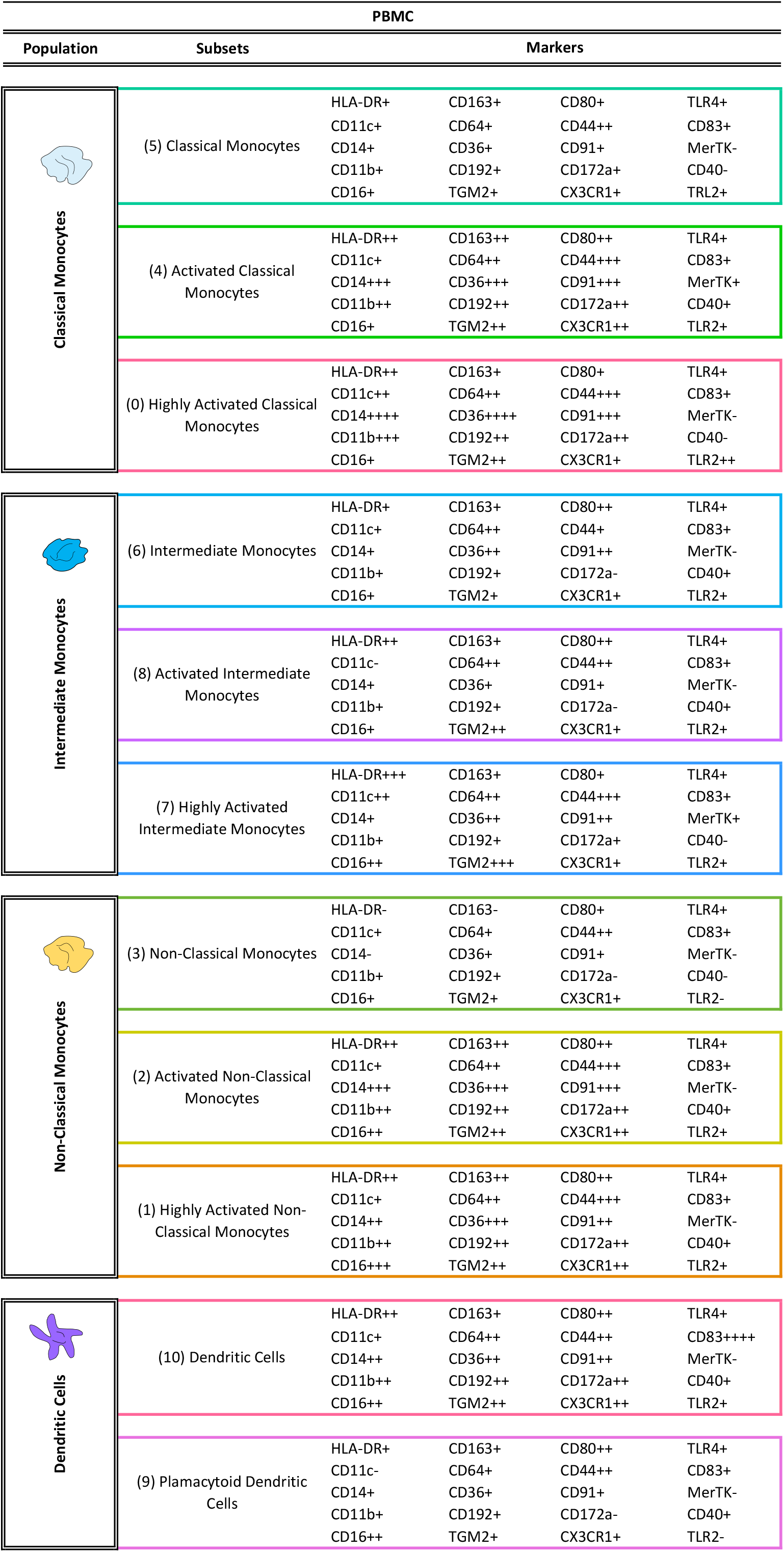
Identified populations in peripheral blood from the unsupervised clustering analysis obtained using the focused panel. The numbers in the parenthesis indicate the number of each cluster showed in the UMAP. Levels of expression were determined by the intensity mean of each marker in each cluster; 0-5 (−), 6-100 (+), 101-500 (++), 501-1000 (+++),> 1001 (++++).

Association analysis performed in the proliferative phase of the menstrual cycle revealed an increased proportion of activated non-classical monocytes and pDCs in blood from women with endometriosis compared to controls and a decrease of classical monocytes, intermediate monocytes, and non-activated non-classical monocytes **(Figure 8. A).** However, significant differences in markers between controls and cases were not found, suggesting that previous differences in marker expression found in endometrium may be tissue specific.

**Figure 8.**
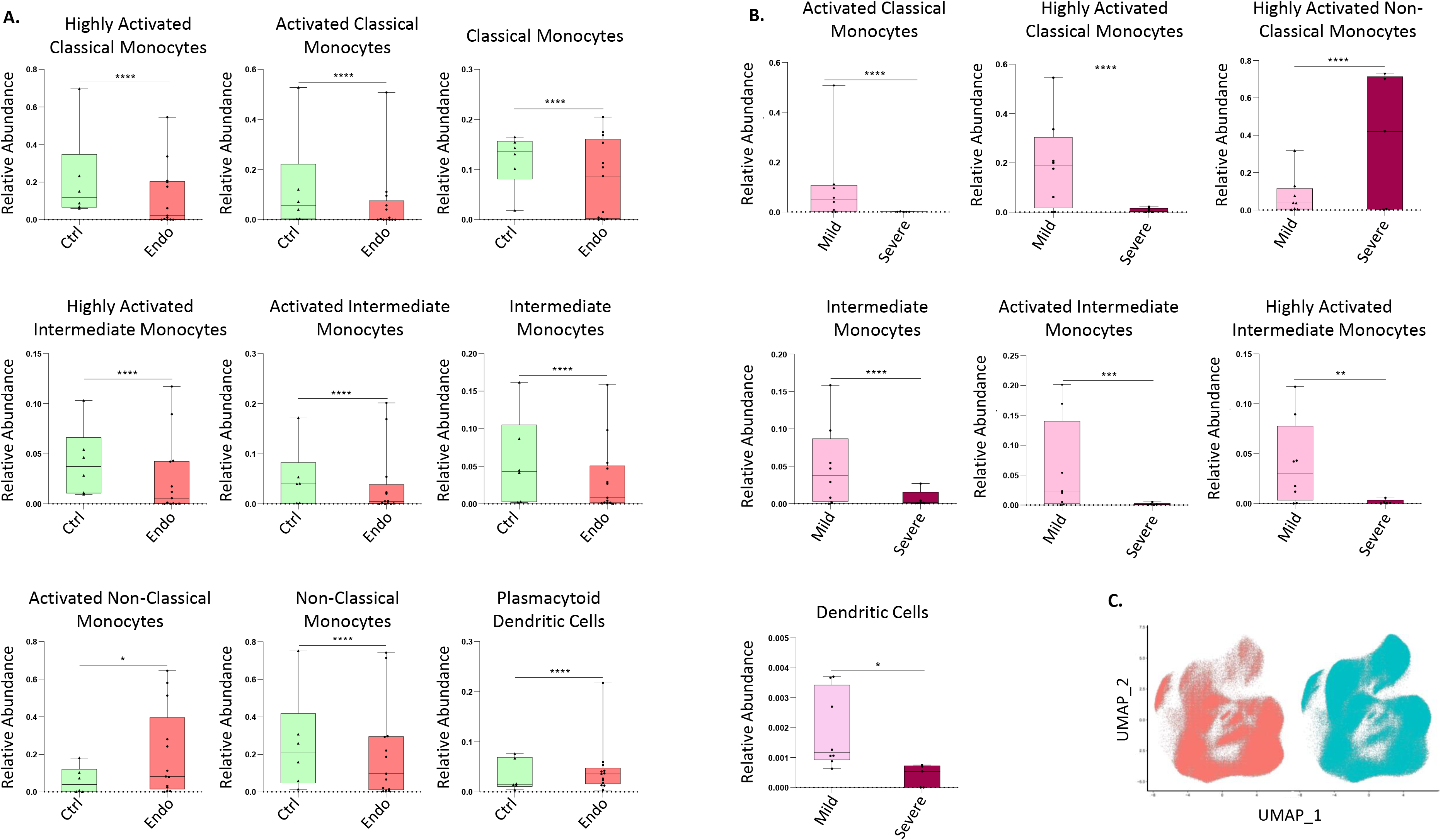
Focused panel association analysis results in blood. **A)** The figure shows the significant differences in abundance of immune populations in the proliferative phase of the menstrual cycle in cases and controls. **B)** Differences in abundance of populations between stages of endometriosis in blood. Analysis performed in the proliferative phase of the menstrual cycle. **C)** UMAP from controls (red) and endometriosis (blue) samples, including both proliferative and secretory phases and both stages of the disease. Arrows show the cluster of highly activated non-classical monocytes. Y axis: Relative abundance of cells per cluster. The total cells per cluster represents all the cells from each specific population from all conditions analyzed (controls, endometriosis, proliferative and secretory phase). Ctrl: controls, Endo: endometriosis. Analysis performed in the proliferative phase of the menstrual cycle. pValues: *, pVal ≤ 0.05; **, pVal ≤ 0.005; **** pVal ≤ 0.00005. n=6 controls and n=13 endometriosis PE (8 mild stage and 5 severe stage). PE: proliferative.

Finally, to evaluate if there could be an association of immune populations in blood with severity of the disease, association analysis of stages of endometriosis in blood were performed. The results showed that classical monocytes (highly activated and activated), intermediate monocytes (intermediate monocytes; activated and highly activated), and dendritic cells are more abundant in peripheral blood from women with mild versus severe disease, suggesting that there is recruitment of monocytes to sites of inflammation in women with endometriosis. In contrast, highly activated non-classical monocytes were significantly higher in the setting of severe disease **(Figure 8. B),** and, interestingly, this population was considerably more abundant in endometriosis than in control patients **(Figure 8. C)**, which could serve as a biomarker of disease and, specifically, severity of endometriosis.

## Discussion

The data herein provide strong support for a compromised innate immune system in endometrium and blood of women with endometriosis. Mass cytometry has revolutionized the cytometry field, allowing simultaneously labeling and analyzing cells of interest at the single cell level with a greater number of antibodies than traditional flow cytometry. To our knowledge, we are the first group to deep phenotype endometrial immune cells using CyTOF, although others have used this technique in peritoneal fluid and blood [21] and in menstrual effluent [22]. Herein, we used two complementary CyTOF panels comprised of 42 and 38 (after exclusion of non-mononuclear phagocytes) markers that enabled deep phenotyping of endometrium and PBMCs from women with and without endometriosis to better understand the role of these populations in disease. To note, previous studies did not use a mononuclear phagocyte-focused panel, which allowed us to identify more detailed differences. Using the broad panel, we identified significantly higher abundance of endometrial macrophages and neutrophils in endometriosis patients compared to controls. We also showed that the dynamics throughout the menstrual cycle of macrophages, dendritic cells, and B cells had contrary direction between both phases in disease and controls. As differences were mainly observed in mononuclear phagocytes, which were also the most abundant among the CD45^+^ cells, we further pursued deep phenotyping of this populations and discovered differences in controls and cases in different hormonal milieu (cycle phases). As the sample size in the secretory phase was small and significant differences were not found in this phase, we focused on the proliferative phase. The latter is estrogen-dominant and is analogous to the initial inflammatory phase for repairing the endometrium after menses, where pro-inflammatory macrophages play a vital role in wound-healing [23]. Thus, this phase provides valuable information to study the immune populations in endometrial function.

### Deep immunophenotyping of mononuclear phagocytes indicates decreased phagocytic capacity of macrophages and dendritic cells in endometrium of women with endometriosis

Our results showing higher abundance of endometrial macrophages in the proliferative phase in women with versus without endometriosis are in accordance with other studies [24]. Also, over-expression of MCP-1 (CCL2) and IL-1β in endometrium of women with endometriosis [25], suggests increased infiltration of monocytes that differentiate to macrophages in this tissue in women with disease **(Figure 9).** Our results show that CD91^+^ macrophages are more abundant and have different dynamics throughout the cycle in cases versus controls. CD91 participates in the efferocytosis of apoptotic cells – a crucial function during endometrial shedding. Notably, we found that endometrial CD91^+^ macrophages and CD1a^+^ dendritic cells over-express SIRPα - which acts to inhibit efferocytosis by these cells - in women with endometriosis compared to controls. Healthy cells express “don’t-eat-me” signals, exposing different markers, such as CD47, that, when exposed, are recognized by SIRPα, and prevent engulfment by initiating a cascade of events leading to the inhibition of phagocytosis. Such engulfment is promoted by the calreticulin/CD91 pathway as an “eat-me” signal. In many cancers, calreticulin is exposed in the cell surface that would indicate an “eat-me” signal. However, this process does not occur in cancer, and why this molecule is exposed in cancerous cells is still unclear (Boada-Romero, *et al.,* 2020). We found that macrophages that overexpress SIRPα in endometriosis are CD91^+^. Even if they express the latter, these observations suggest that their phagocytic capacity may be compromised by concomitant overexpression of SIRPα, suggesting that endometrial cells in women with endometriosis behave similarly to cancer cells, sending signals that interfere with the CD91 pathway in macrophages preventing efferocytosis. Thus, the overexpression of this marker in specific myeloid cells may be a consequence of aberrant functionality of eutopic endometrial cells in women with endometriosis. Xie *et al.,* demonstrated that *in vitro* cultured macrophages treated with endometrial homogenates from women with endometriosis had significantly increased SIRPα expression, not observed when the macrophages were treated with homogenates of endometrium from women without disease [27], also suggesting that signals from eutopic endometrium affect the phagocytic capacity of macrophages. Another study evaluated the effect of this pathway in endometrial stromal cells from endometrium of women with endometriosis and in endometriosis lesions *in vitro* [28]. Co-culture of PBMCs and lesion-derived stromal cells resulted in up-regulation of SIRPα in macrophages, further supporting paracrine endometrial cell interactions in macrophage dysfunction and disease establishment. Notably, we found that endometrial CD1a^+^ dendritic cells also overexpress SIRPα in women with versus those without endometriosis, indicating that other endometrial phagocytic cells may also be dysregulated in women with endometriosis. Decreased phagocytic capacity in macrophages and dendritic cells may allow aberrant endometrial cells to escape immunologic surveillance during tissue shedding, leading to establishing pelvic endometriosis **(Figure 9).** Sampson’s theory of the origin of endometriosis proposes that endometrial cells shed during menses survive and migrate to the peritoneal cavity through retrograde menstruation, where they implant and develop endometriotic lesions [29]. Our results, together with the above studies, suggest that endometrial macrophages have defective phagocytic capacity in women with versus without endometriosis, opening opportunities for mining new targets to prevent development and/or progression of the disease as well as endometrial dysfunction. Expression of calreticulin in endometrial cells in women with endometriosis and the effects in the calreticulin/CD91 pathway warrant further study to confirm this paradigm.

**Figure 9.**
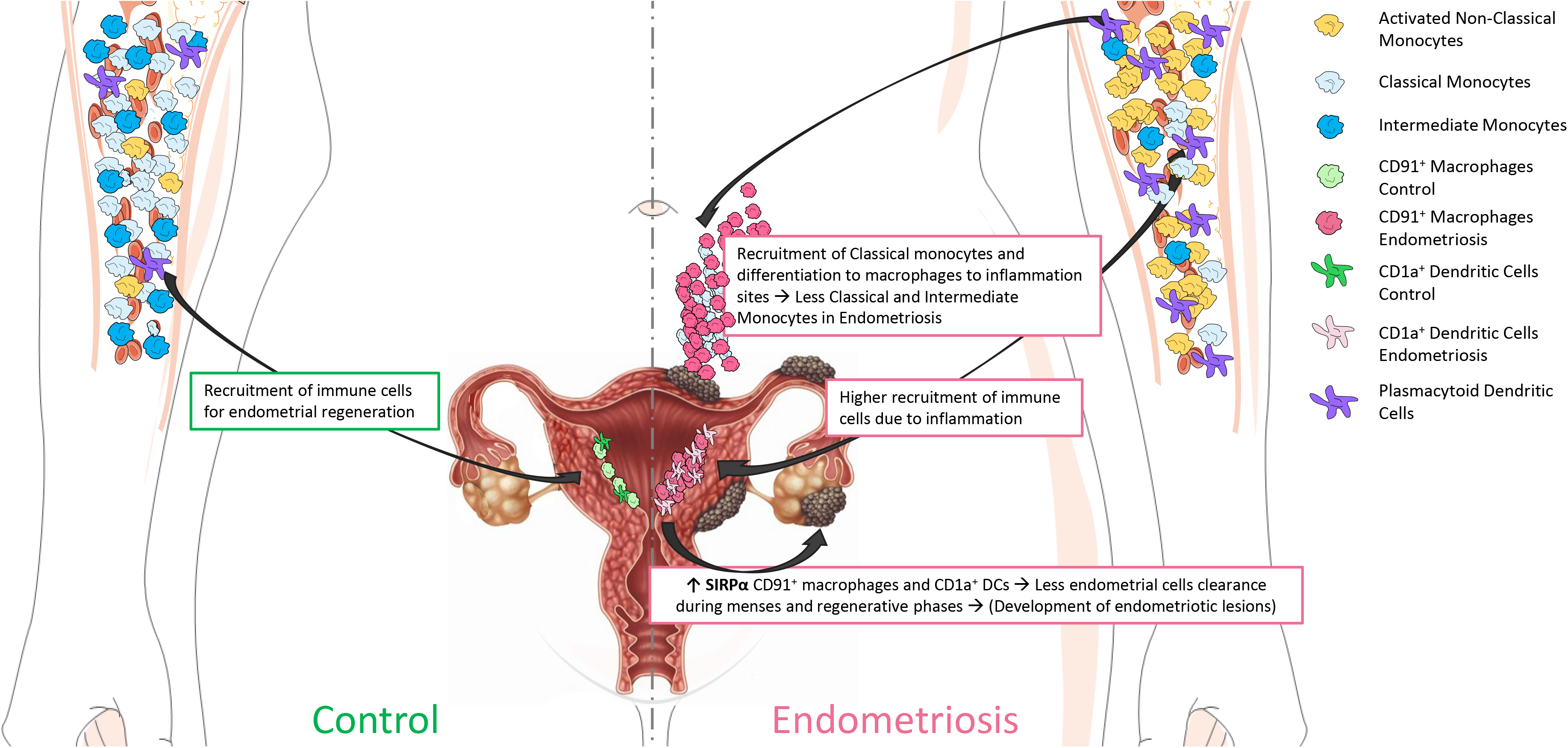
Proposed model of involvement of specific immune populations in the proliferative phase of the menstrual cycle in endometriosis patients compared to controls. The model shows that, in controls, there is a recruitment of immune cells during regenerative-proliferative phase of the menstrual cycle for endometrial regeneration and proliferation. In addition, there is less recruitment of classical and intermediate monocytes to the endometrium in controls, as the inflammation is not as enhanced in this tissue as it is in disease. The right panel shows the model in endometriosis. It shows that there is higher recruitment of immune populations (specifically classical and intermediate monocytes, which will differentiate to macrophages) to sites of inflammation, such as the endometriotic lesions and endometrium, decreasing like this the proportion of these monocytes in circulation of women with endometriosis. It also shows that there is a higher proportion of plasmacytoid dendritic cells and non-classical monocytes in circulation in women with endometriosis. Finally, we hypothesize that the increase of SIRPα in endometrial macrophages and dendritic cells might decrease the phagocytic capacity of these cells, by allowing endometrial cells to scape clearance during menses and regenerative phases of the cycle, which, in turn, would allow their migration to the peritoneal cavity, implant and develop the endometriotic lesions.

In addition, our findings that CD91^+^ endometrial macrophages are more abundant and overexpress CD64 in women with endometriosis and in mild versus severe stage disease suggest a more enhanced pro-inflammatory phenotype in mild disease due to the expression of CD64. Therapeutic targeting of CD64 expressed on pro-inflammatory macrophages, proposed for other chronic inflammatory diseases, may be useful in the context of endometriosis [30].

### Immunophenotyping suggests higher infiltration of circulating monocytes to sites of inflammation in women with endometriosis

Discovery herein has revealed that in addition to endometrial immune populations, specific PBMC populations differ in women with versus without endometriosis. Importantly, classical and intermediate monocytes are in lower abundance and pDCs and activated non-classical monocytes are more abundant in blood from women with versus without disease. These findings are in line with a recent study that described decreased phagocytic function of monocytes in blood from patients with endometriosis before surgery, and normalization after lesion removal, compared to controls undergoing surgery but without endometriosis [31]. In addition, proliferation of endometrial cells *in vitro* from women with endometriosis is significantly enhanced by blood monocytes; whereas, monocytes from healthy patients inhibit their proliferation [32], indicating factors influencing these cells act as enhancers or suppressors of endometriotic lesion development. Our results that classical and intermediate monocytes are less abundant in blood from patients with endometriosis are consistent with increased recruitment of monocytes to sites of inflammation, e.g., to the peritoneal cavity, where endometriotic lesions are commonly found, and to the uterine endometrium, where we also observed a higher proportion of macrophages in women with disease. In addition, these populations may have less phagocytic function and thus play a role in deficient clearance of ectopic endometrial tissue, resulting in endometriotic lesion development **(Figure 9).** While the above studies by others did not differentiate findings in the setting of mild and severe disease or different types of monocytes, our results show that this effect is more profound in severe disease, where there are significantly less circulating classical and intermediate monocytes than in mild disease, suggesting this defective function is enhanced in more advanced disease.

### Abundance of specific mononuclear phagocytes in blood are putative biomarkers of disease and severity of endometriosis

Notably, we found an increase of activated non-classical monocytes and pDCs in blood of women with endometriosis versus controls and that the former were more associated with advanced stage disease. Suen *et al.,* demonstrated in a murine model that IL-10 secreted from pDCs promotes endometriotic lesion development through aberrant angiogenesis early in disease establishment [33]. Our results show that pDCs are associated with disease but not stage. This is in contrast to the mouse model, where the association with early disease establishment can be assessed experimentally, unlike human endometriosis where the evolution cannot be reliably assessed. It is well known that pDCs are involved in some autoimmune diseases and play a pivotal role in the development of autoantibodies through impaired Type I interferon (Type I INF) production [34], enhancing systemic autoimmunity [35]. Endometriosis is a chronic inflammatory disease that shares many similarities with autoimmune disorders [36], including production of autoantibodies [37]. A recent metanalysis by Shigesi *et al.,* found a greater risk of autoimmune diseases in patients with endometriosis, including systemic lupus erythematosus, Sjögren’s syndrome, rheumatoid arthritis, celiac disease, multiple sclerosis, inflammatory bowel disease, autoimmune thyroid disorders, and Addison’s disease [38]. In addition, more severe endometriosis is being associated with concomitant autoimmune diseases [39]. Thus, it is of great interest to determine the precise role of pDCs in endometriosis pathophysiology and the role of systemic cytokines, such INF family members, to better understand common immunopathology between endometriosis and autoimmune diseases. Interestingly, in the autoimmune disorder neuromyelitis optica spectrum disorder (NMOSD), non-classical monocytes have a higher frequency than in controls in blood [40]. Non-classical monocytes are believed to be patrolling and anti-inflammatory innate immune cells. However, their roles in chronic diseases, such as in multiple sclerosis or systemic lupus erythematosus, are less clear as their functions can be protective as well as positively associated with disease burden [41]. Given our observation of their greater abundance in blood of women with endometriosis and, specifically, in those with severe disease, and as endometriosis is a chronic inflammatory disorder, further study of the functional roles of this population and mining biomarkers of disease severity warrants further investigation. Although the patients in the current study did not have autoimmune comorbidities, further analysis of abundance and function of non-classical monocytes and pDCs, as well as their cytokine production in patients with endometriosis, having or not concomitant autoimmune diseases, is of interest, especially as the role of these populations in endometriosis remains unresolved.

Some of the limitations of the study include a limited sample size, especially in the secretory phase of the menstrual cycle. As mentioned above, this may be a reason why we found only differences in the proliferative phase. Although in this study we show differences in the latter, a larger sample size in all phases would further confirm differences highlighted by our discovery cohort. Another limitation of the study is that the focused panel did not contain a B cell-specific marker. As we observed differences in B cell frequency by using the broad panel, indication that B cells may also have a role in disease would require further investigation. Along the same line, while we were able to identify coarsely dendritic cell populations, additional markers deeply characterizing them would be needed to distinguish between known subsets such as cDC1 and cDC2. Although we had some limitations in the study, to our knowledge, this cohort size represents the largest cohort of endometrial samples from women with and without endometriosis analyzed by mass cytometry reported to date. Applying two different panels enabled us to identify populations that could be involved in endometriosis, and the second panel allowed deep phenotyping the MPC system at the protein level for the first time in eutopic endometrium of women with and without endometriosis. Considering the heterogeneity of the disease and the immune complexity, further single cell studies, e.g., using more focused panels for other immune populations and endometrial cells, will help to better understand the immune system in endometriosis.

## Conclusions

The data presented herein support that endometrial CD91^+^ macrophages and CD1a+ dendritic cells of women with endometriosis display a decreased phagocytic capacity compared to controls. If this is a consequence of aberrant endometrial cell functions or a defect on the immune system, per se, awaits further investigation. In addition, classical and intermediate monocytes are less abundant in blood from women with endometriosis, suggesting higher recruitment of these populations to sites of inflammation and disease lesions, with current evidence demonstrating defective phagocytic function in the tissues and in the circulation. Finally, pDCs and activated non-classical monocytes are more abundant in endometriosis, with the latter being even higher in severe disease, which could serve as a biomarker of disease severity in blood. Overall, we conclude that the mononuclear phagocyte system is compromised in endometrium and peripheral blood of patients with endometriosis. The results of our study open alternative venues for developing new diagnostic and therapeutic targets and strategies for identifying and treating sub-types of this enigmatic disorder.

## Supporting information

Supplemental Figure 1. Manual gating strategy for the focused panel. The figure shows an example of the dot plots obtained using FlowJo in endometrial

Supplemental Figure 2. Endometrial sampling methods. The figure shows UMAPs of the two methods used for endometrial tissue collection, biopsy (red) an

Supplemental Figure 3. Batch effects derived from different runs. The figure shows UMAPs representing the distribution of the cells from the runs in t

Supplemental Figure 4. Proportion of endometrial immune populations identified by using the broad panel. The figure shows the proportion of each popul

Supplemental Figure 5. Fluctuation of all endometrial immune populations identified in the broad panel. A) Differences in abundance of populations th

Supplemental Figure 6. Fluctuation of all endometrial immune populations identified in the focused panel. A) Differences in abundance of populations

Supplemental Table 1. Samples used in the study. Endo: endometriosis; PE: proliferative; SE: secretory: MSE: mid secretory, LSE: late secretory; N/A:

Supplemental Table 2. CyTOF panels used in the study. Left: broad panel, right: focused panel.

Supplemental Table 3. Expression of the significant markers between endometriosis and controls in all cell types in endometriosis versus controls from

## List of abbreviations

ADC: Acid citrate dextrose
BH: Benjamini-Hochberg
BSA: Bovine serum albumin
Ctrl: Control
CyTOF: Cytometry by-time-of-flight
EDTA: Ethylenediaminetetraacetic acid
Endo: Endometriosis
ESE: Early secretory
FACS: Fluorescent activated cell sorting
FDR: False discovery rate
GLMM: Generalized linear mixed model
INF: Interferon
LSE: Late secretory
MPC: mononuclear phagocytic cells
MSE: Mid secretory
MSI: Mean signal intensity
PBMCs: Peripheral blood mononuclear cells
PE: Proliferative
PFA: Paraformaldehyde
Rpm: Revolutions per minute
RT: Room temperature
SCM: Serum containing media
SE: Secretory
SNN: Shared nearest neighbor
TCEP: Tris (2-carboxyethyl) phosphine hydrochloride

## Declarations

### Ethics approval and consent to participate

Specimens were obtained from the UCSF/NIH Human Endometrial Tissue Bank under IRB approval (IRB#10-02786) from 2019 to 2021. Written informed consent was obtained from all participants.

### Consent for publication

“Not applicable”.

### Availability of data and materials

The datasets used and/or analyzed during the current study are available from the corresponding author on reasonable request.

### Authors’ contributions

<u>JVJ:</u> processing of samples, design of panels, acquisition, analysis, and interpretation of data, drafting, and reviewing the article; <u>SS:</u> processing of samples and acquisition of data; <u>RT:</u> data analysis, drafting, and reviewing the article; <u>MGS:</u> data analysis, drafting, and reviewing the article; <u>DK:</u> processing of samples, design of panels, acquisition of data; <u>JV:</u> design of panels and reviewing the article; <u>AFG:</u> experimental design and optimization of protocols, and reviewing the article; <u>KCV:</u> collection of samples; <u>JCI:</u> interpretation of data and reviewing the article; <u>NRR:</u> experimental design, optimization of protocols, and reviewing the article; <u>AJC:</u> design of panels, analysis, and interpretation of data, drafting, and reviewing the article; <u>LCG:</u> experimental design, data interpretation, drafting, writing, and reviewing and editing the article.

## Acknowledgements

We acknowledge Tristan Coureau, Nayvin Chew, Stanley Tamaki, Bushra Samad, Gabriella Fradgiadakis, for their help in teaching some of the used protocols. We also acknowledge the NIH S10 1S10OD021822-01 grant from the UCSF Flow CoLab.

