## Supplementary figures and images for "Deep Immunophenotyping Reveals Endometriosis is Marked by Dysregulation of the Mononuclear Phagocytic System in Endometrium and Peripheral Blood"

### Supplemental Figure 1. Manual gating strategy for the focused panel. The figure shows an example of the dot plots obtained using FlowJo in endometrial

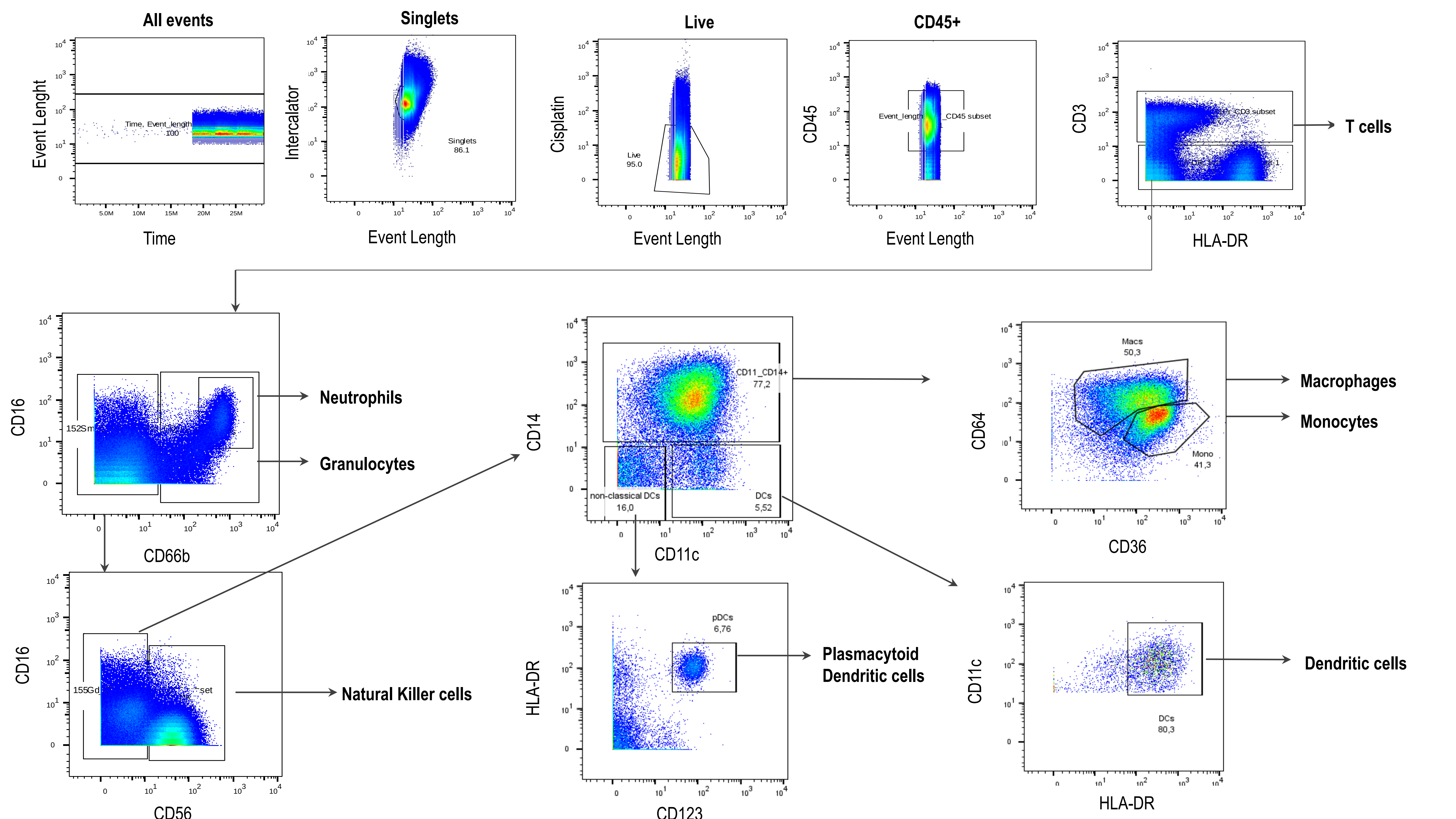

### Supplemental Figure 2. Endometrial sampling methods. The figure shows UMAPs of the two methods used for endometrial tissue collection, biopsy (red) an

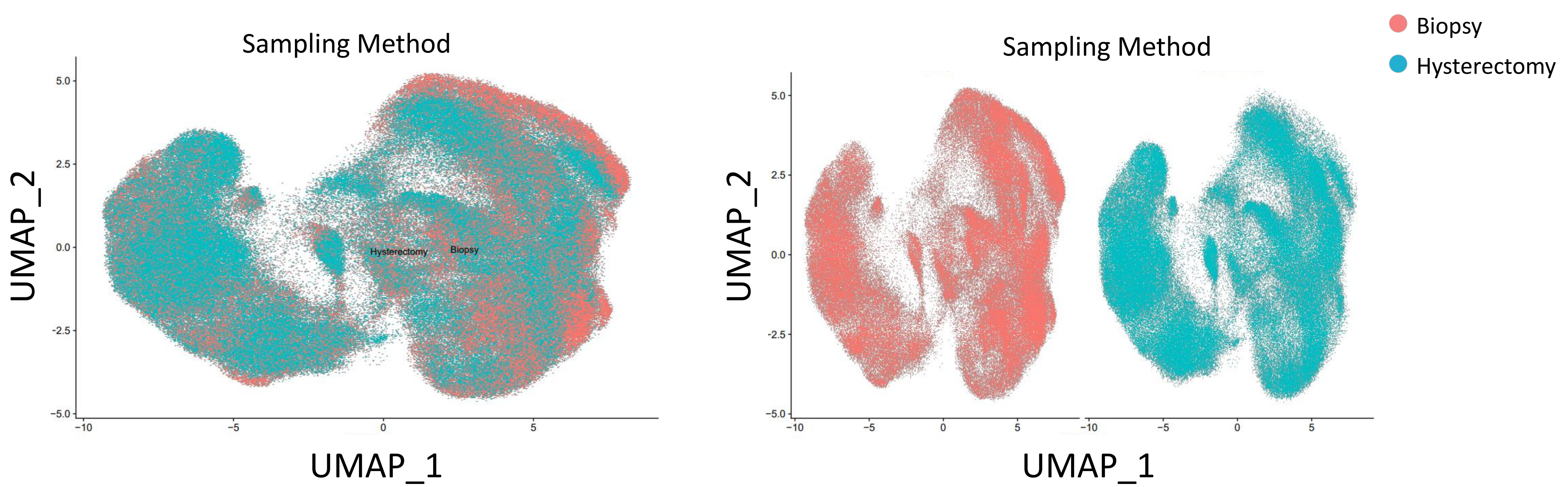

### Supplemental Figure 3. Batch effects derived from different runs. The figure shows UMAPs representing the distribution of the cells from the runs in t

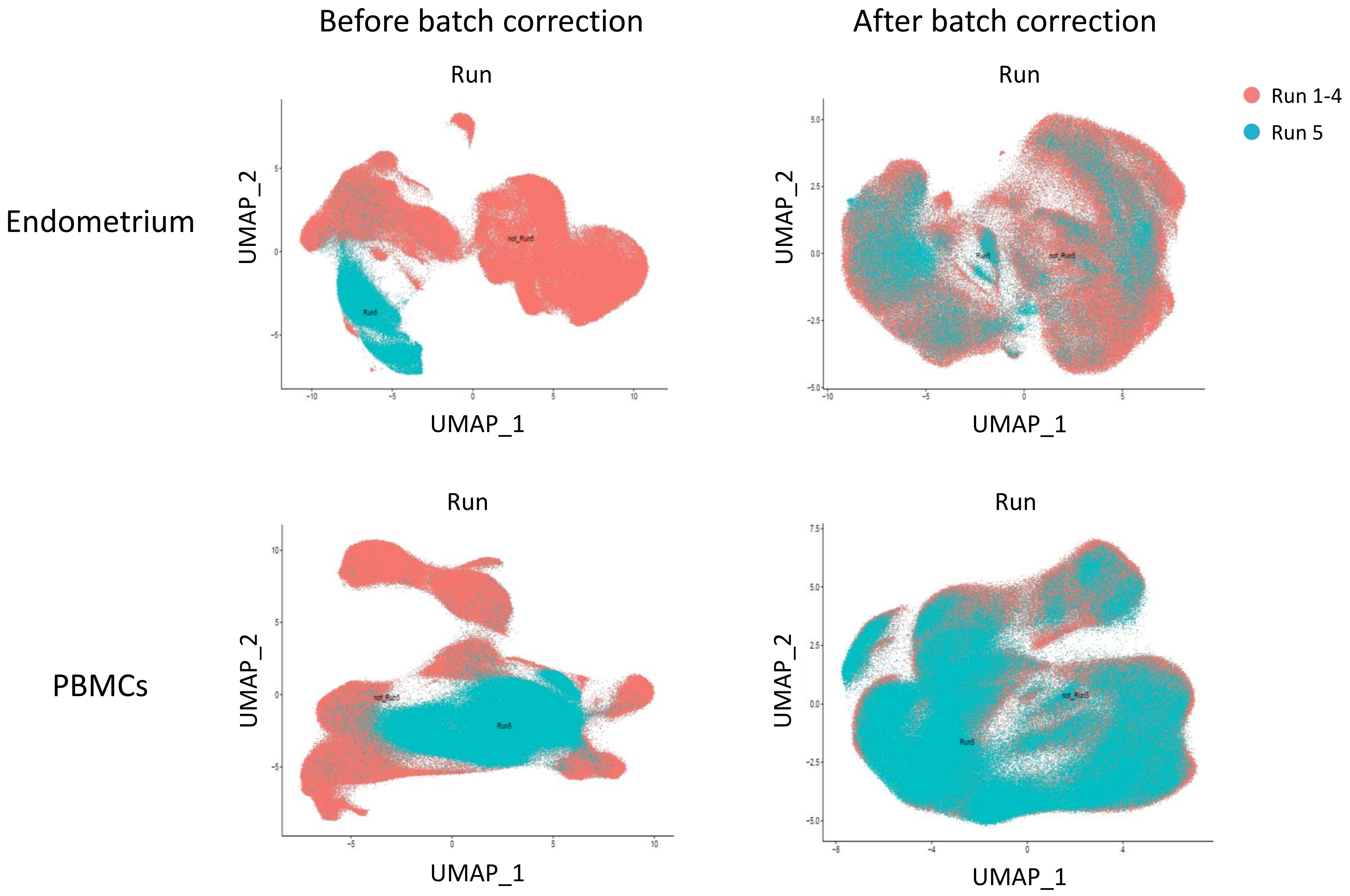

### Supplemental Figure 4. Proportion of endometrial immune populations identified by using the broad panel. The figure shows the proportion of each popul

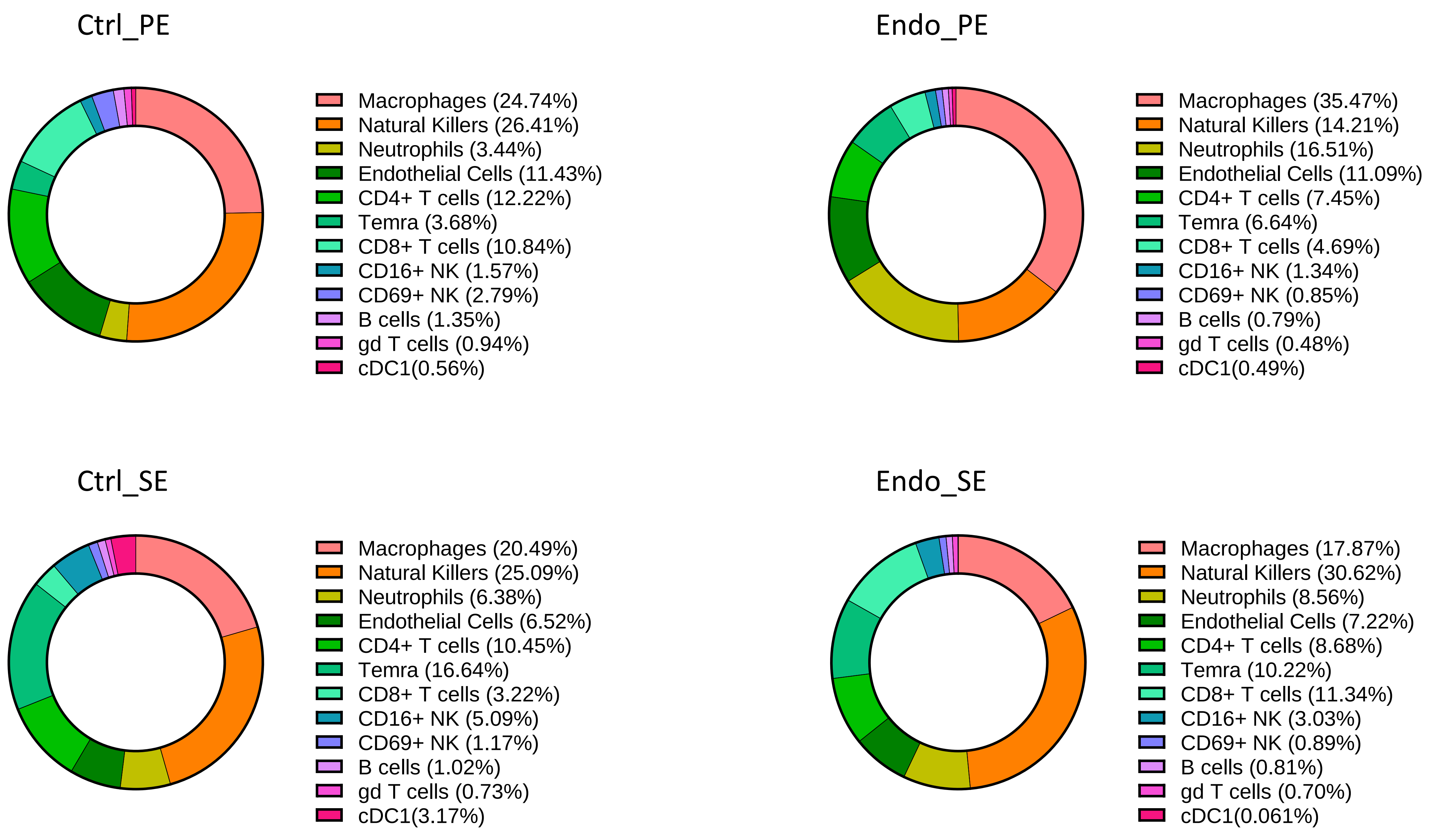

### Supplemental Figure 5. Fluctuation of all endometrial immune populations identified in the broad panel. A) Differences in abundance of populations th

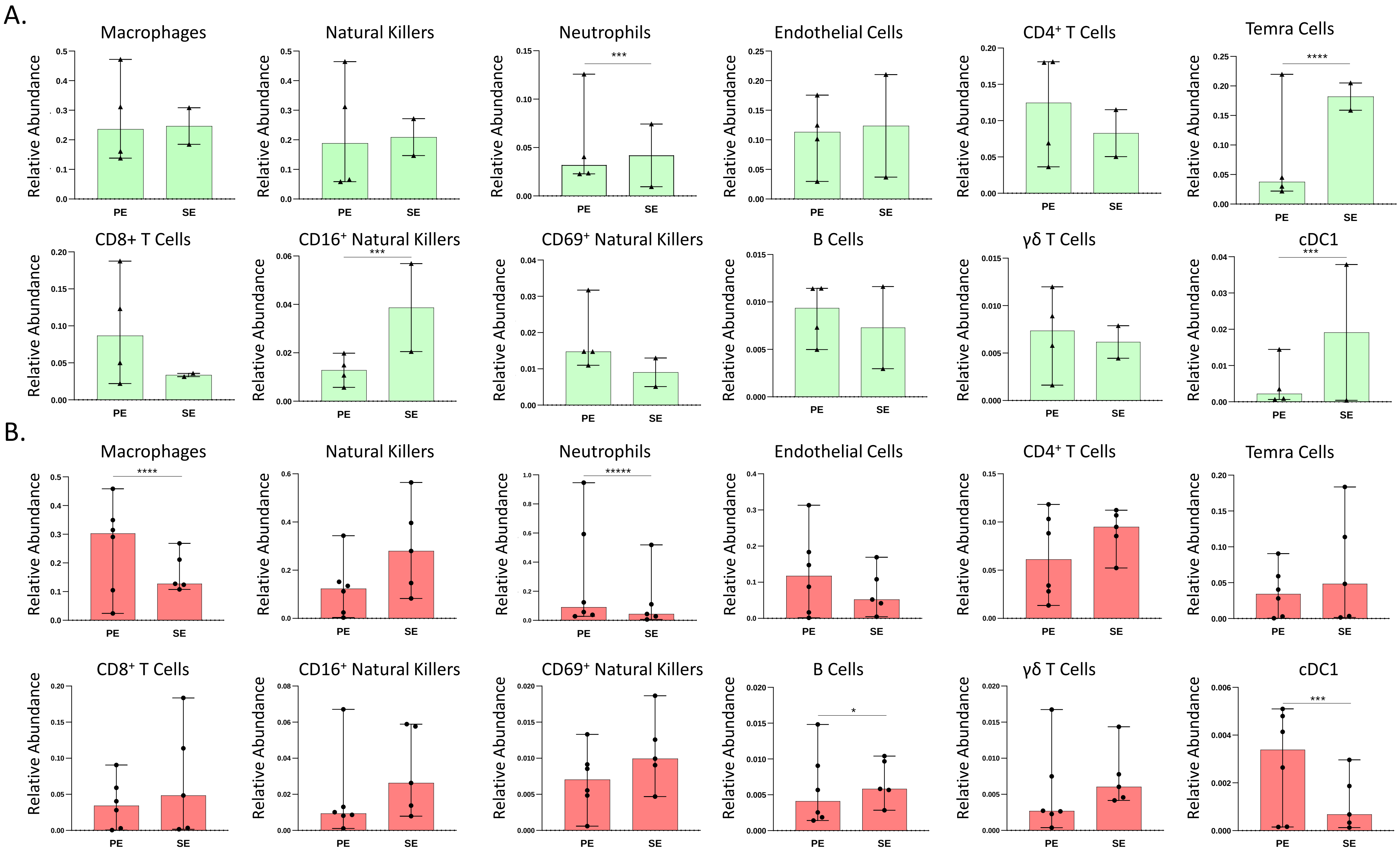

### Supplemental Figure 6. Fluctuation of all endometrial immune populations identified in the focused panel. A) Differences in abundance of populations

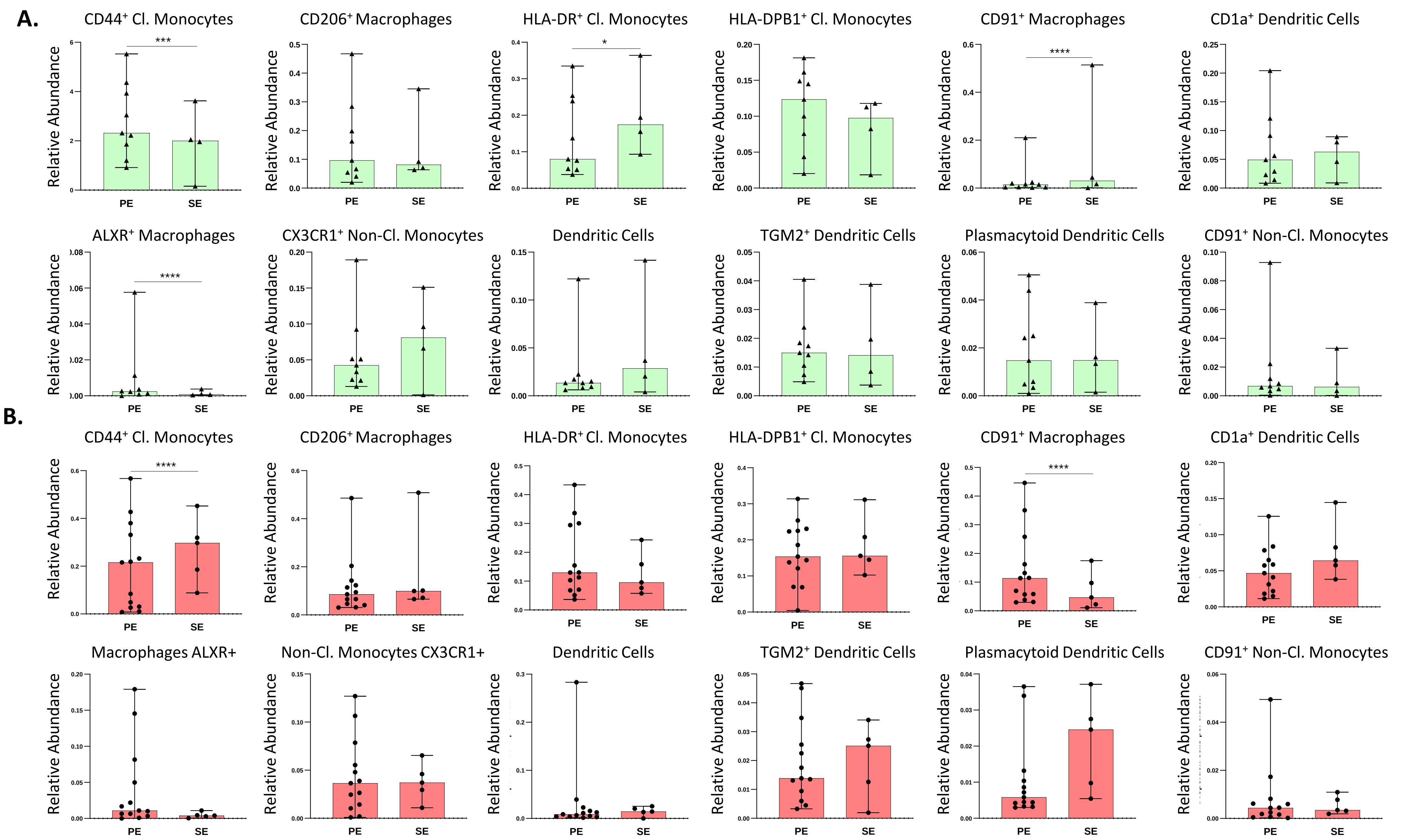
